# Logic-based machine learning predicts how escitalopram attenuates cardiomyocyte hypertrophy

**DOI:** 10.1101/2024.11.13.623416

**Authors:** Taylor G. Eggertsen, Joshua G. Travers, Elizabeth J. Hardy, Matthew J. Wolf, Timothy A. McKinsey, Jeffrey J. Saucerman

## Abstract

**Introduction:** Cardiomyocyte hypertrophy is a key clinical predictor of heart failure. High-throughput and AI-driven screens have potential to identify drugs and downstream pathways that modulate cardiomyocyte hypertrophy.

**Methods:** Here we developed LogiRx, a logic-based mechanistic machine learning method that predicts drug-induced pathways. We applied LogiRx to discover how drugs discovered in a previous compound screen attenuate cardiomyocyte hypertrophy. We experimentally validated LogiRx predictions in neonatal cardiomyocytes, adult mice, and two patient databases.

**Results:** Using LogiRx, we predicted anti-hypertrophic pathways for 7 drugs currently used to treat non-cardiac disease. We experimentally validated that escitalopram (Lexapro) and mifepristone inhibit hypertrophy of cultured cardiomyocytes in two contexts. The LogiRx model predicted that escitalopram prevents hypertrophy through an “off-target” serotonin receptor/PI3Kγ pathway, mechanistically validated using additional investigational drugs. Further, escitalopram reduced cardiomyocyte hypertrophy in a mouse model of hypertrophy and fibrosis. Finally, mining of both FDA and University of Virginia databases showed that patients with depression on escitalopram have a lower incidence of cardiac hypertrophy than those prescribed other serotonin reuptake inhibitors that do not target the serotonin receptor.

**Conclusion:** Mechanistic machine learning by LogiRx discovers drug pathways that perturb cell states, which may enable repurposing of escitalopram and other drugs to limit cardiac remodeling through “off-target” pathways.

## INTRODUCTION

Cardiac hypertrophy is a key clinical predictor of heart failure^1–5^. Hypertrophy is a response to various physiological stressors, such as pressure overload, drug toxicity, or gene mutations^6^. Several drugs prescribed for heart failure also affect cardiac hypertrophy, such as angiotensin converting enzyme (ACE) inhibitors, beta blockers, and angiotensin receptor blockers (ARBs)^7^. But heart failure rates are still rising, with a five year mortality rate > 40%^8^. There has been increasing effort towards drug repurposing for cardiovascular indications to accelerate translation and reduce cost^9,10^. One such success is the repurposing of SGLT2 inhibitors from diabetes to cardiovascular disease^11^.

Numerous medications not initially designed for cardiac treatment impact the progression of cardiac hypertrophy^12–15^, although their mechanisms are largely unknown. Experimental approaches to prioritize drugs from large screens and investigate their mechanisms are time intensive and costly. Drug development is further hindered by the complexity of hypertrophic signaling, with a high degree of cross-talk between multiple molecular pathways^7,16^. Protein interaction databases and gene regulatory networks have been used previously to explore drug target interactions^17,18^, however these studies have primarily examined network topology rather than simulating network response to drugs.

To address the challenges of mapping and mechanistically understanding drug responses from phenotypic screens, we developed a logic-based predictor of drug pathways (LogiRx). LogiRx maps from drug-target and protein interaction databases to curated logic-based network models of cell phenotype. We used LogiRx to prioritize seven anti-hypertrophic drugs identified in a previous cell-based screen^12^, providing mechanistic insight to how two drugs attenuate hypertrophy via “off-target” pathways. LogiRx predictions were experimentally validated in cultured cardiomyocytes, mice, and in two patient databases of electronic health records. Together, these studies validate the LogiRx method and provide mechanistic insights into how escitalopram (Lexapro) and other drugs modulate cardiomyocyte hypertrophy.

## METHODS

### Linking drug targets to logic-based signaling network using LogiRx

We used a published logic-based differential equation model of the cardiomyocyte hypertrophy signaling network^19^. This literature-based network of proteins and genes is composed of 107 nodes (proteins or mRNAs) and 193 edges, validated against 450 experiments across the literature with 77% accuracy^20^. The model correctly predicted the anti-hypertrophic effect of 28 out of 32 drug responses from our previous experimental screen^12^.

### Linking drug targets to logic-based signaling network using LogiRx

We first identified hits from a previous screen for compounds that inhibit cardiomyocyte hypertrophy^12^, identifying protein targets using the drug-target database DrugBank^21^ (**Figure 1A**). LogiRx identified top scoring directed pathways from drug targets to the hypertrophy signaling network nodes through Omnipath directed interactions^22,23^ using the PathLinker algorithm^24,25^ in Cytoscape^26^ (**Figure 1B**). Top scoring candidate drug pathways were added to a LogiRx-expanded logic-based network model using Netflux.

**Figure 1.**
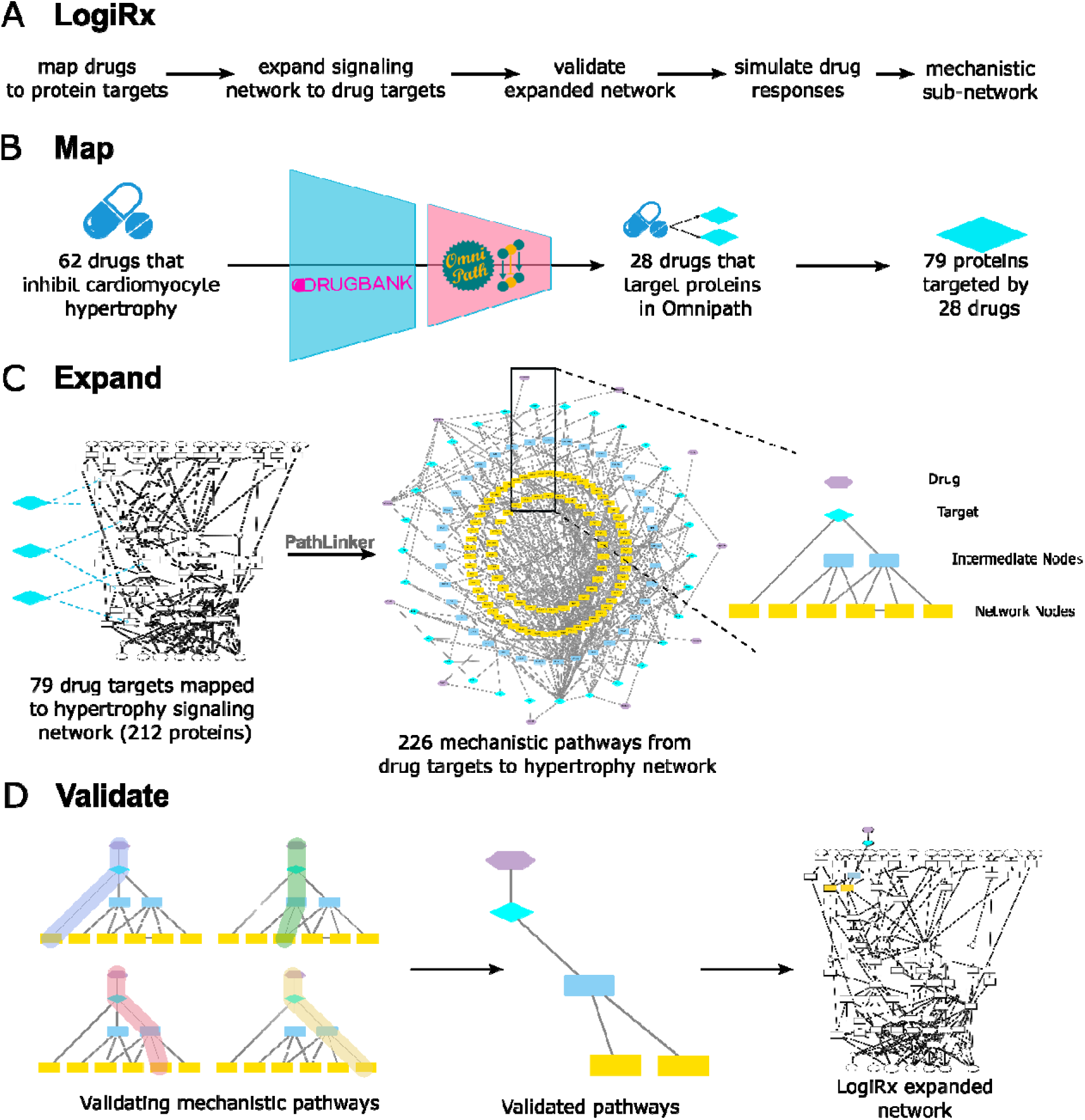
LogiRx discovers pathways from anti-hypertrophic drugs to the cardiomyocyte hypertrophy signaling network. A) Overview of the LogiRx workflow, from drugs to mechanistic predictions. B) Drug targets are identified from DrugBank and filtered by annotations in the protein-interaction database Omnipath. C) Drugs and their protein targets are mapped to the hypertrophy network through LogiRx. D) Drug pathways are individually simulated using Netflux and validated against prior experimental literature. Drug pathways that maintain model validation accuracy are used to expand the hypertrophy network for drug predictions.

### Literature validation of LogiRx-expanded models

The LogiRx-expanded logic-based differential equation model was simulated to test the functional impact of the drug pathways on cardiomyocyte hypertrophy. Each pathway was individually added to the signaling network model to create expanded logic-based differential equation model variants. Each drug-pathway-expanded model was simulated to predict the semi-quantitative network dynamics in response to drug. To assess the functional validity of the LogiRx-identified drug pathways, we validated their performance against 450 literature experiments previously used to validate the hypertrophy signaling model^20^ (**Figure 1C**).

### Simulating drug mechanisms of action

Drug mechanisms of action were implemented using logic-based equations as described previously^27,28^, accounting for mechanisms of drug action (agonist or antagonist) and drug binding (competitive or non-competitive). The binding properties determine how the upstream signal is shifted by the drug dose, while the drug action determines whether the target node is upregulated (agonist) or downregulated (antagonist). Each simulation was run first without drug input to establish baseline activity, and then with the appropriate drug to identify steady-state changes in drug response.

### Mechanistic subnetwork analysis

To identify pathways mediating drug response, we utilized the mechanistic subnetwork method^27–29^. We simulated the global response of the network to the drug in the presence of hypertrophic stimulus to identify “drug-responsive nodes”. We then simulated knockdown of each node and measured that node’s influence on drug-induced change in cardiomyocyte hypertrophy, referring to these nodes as “regulating nodes”. The intersection of the regulating nodes with responsive nodes results in a mechanistic subnetwork that maps the pathways mediating drug activity. Thus, LogiRx combines drug-target and protein interaction databases, path optimization, and logic-based network simulations to obtain mechanistic subnetworks that explain drug response.

### Experimental validation in cardiomyocytes

We isolated neonatal cardiomyocytes using the NeoMyts kit from Cellutron as described previously^28^. Cardiomyocytes were cultured with serum for 24 hr in 96-well microplates, followed by a 24 hr serum starve. Cardiomyocytes were then treated with one of two hypertrophic stimuli (10 μM phenylephrine, 5 ng/ml transforming growth factor β), 10% FBS (positive control), or serum-free media alone (negative control). At the same time, cells were treated with specified concentrations of the appropriate compound (escitalopram, mifepristone, paroxetine, sarpogrelate hydrochloride, or AL082D06) for 48 hrs.

To prepare for immunofluorescent imaging, cardiomyocytes were first fixed with 4% paraformaldehyde for 20 min, and then permeabilized with 0.1% Triton-X for 15 min. Cardiomyocytes were blocked with 1% bovine serum albumin in PBS for 1 hr, then treated with mouse anti-α-actinin primary antibody (Sigma-Aldrich Cat#A7811, RRID:AB_476766) at a concentration of 1:200 overnight. Cardiomyocytes were blocked with 5% goat serum in PBS for 1 hr, then Alexa Fluor-568-conjugated goat anti-mouse secondary antibody (Thermo Fisher Scientific Cat#A11031, RRID:AB_144696) at a concentration of 1:200 was applied for 1 hr. The cells were stained with DAPI prior to imaging.

High-content imaging was performed on the stained cardiomyocytes using an Operetta CLS High Content Analysis System. These images were processed using CellProfiler^30^ using a cellular segmentation algorithm developed previously and validated to within 5% of two independent manual segmentations^31^. Median cell area was used as a representative measure of the cell population in each well, and cells with undetectable cytoplasm were not counted.

Total protein was isolated from neonatal rat ventricular myocytes (NRVM) using RIPA lysis buffer (25 mM Tris pH 7.6, 150 mM NaCl, 1% NP-40, 1% sodium deoxycholate and 0.1% SDS) supplemented with Halt™ protease and phosphatase inhibitor cocktail (ThermoFisher 78442). Proteins were separated using SDS-PAGE and transferred onto nitrocellulose membrane (0.45 μm, Thermo Scientific 88018). Total protein was first evaluated using LI-COR Revert™ 520 Total Protein Stain (926-10021), followed by blotting with antibodies directed against phosphorylated protein kinase D (1:1,000; CST 2051) and glyceraldehyde phosphate dehydrogenase (1:5,000; Proteintech 60004-1-Ig) overnight at 4°C in intercept® (TBS) Blocking Buffer (LI-COR 927-60001) containing 0.2% Tween-20. Following incubation with primary antibodies, blots were rinsed and probed with appropriate IRDye® secondary antibodies (LI-COR). Protein bands were visualized using a LI-COR Odyssey XF scanner.

### Angiotensin II / Phenylephrine Infusion in Mice

All animal procedures were performed in accordance with the Institutional Animal Care and Use Committee at the University of Colorado Anschutz Medical Campus. Ten-week-old male C57BL/6J were purchased from The Jackson Laboratory (Strain # 000664). Alzet mini-osmotic pumps (Model 2004), or mock pumps for Sham animals, were implanted in a subcutaneous pocket created at the suprascapular region under isoflurane anesthesia. Osmotic pumps were prepared to deliver 1.5 μg/kg/day Angiotensin II (Bachem H-1705.0100) and 50 μg/g/day (R)-(-)-Phenylephrine hydrochloride (Sigma P6126) to induce systemic hypertension, cardiac hypertrophy and fibrosis. Beginning 24 hours post-osmotic pump implantation, animals were randomly divided into groups receiving 10 mg/kg/day escitalopram oxalate (Selleckchem S4064) in 10% DMSO dissolved in 10% (2-hydroxypropyl) β-cyclodextrin (Sigma H107) in water, or vehicle control, for 14 days.

### Echocardiographic Assessment of Cardiac Structure and Function

Serial transthoracic echocardiographic and Doppler analysis were performed using the VisualSonics Vevo F2 instrument. Animals were anesthetized with 2% isoflurane, hair on the chest was removed using a chemical depilatory, and body temperature maintained at 37°C. Isoflurane was maintained at 1.5% throughout the procedure to ensure a consistent plane of anesthesia across subjects. Parasternal long axis (PSLAX) and parasternal short axis (SAX) views of the left ventricle were obtained; SAX two-dimensional views of the LV at the level of the papillary muscle were used to acquire M-mode images. Anterior and posterior LV wall thickness and internal diameters were measured in systole and diastole using these M-mode images. Mitral inflow Doppler signals and myocardial tissue movement of the mitral annulus were obtained to calculate the ratio of early and active filling waves of blood flow through the mitral valve and ventricular tissue velocity to assess cardiac diastolic function. Speckle-tracking strain analyses were acquired from PSLAX B-mode videos. All echocardiographic measurements and analyses were averaged from at least four cardiac cycles and analyses were performed in a blinded manner by a dedicated small animal echocardiography team.

### Tissue Procurement

At study endpoint, animals were sacrificed by exsanguination and hearts excised and placed in ice-cold saline. Right ventricular (RV) tissue was dissected from the LV by cutting along the septum and the outer wall of the LV, and all parts weighed. 50 mg base and apical biopsies of the LV were flash-frozen in liquid nitrogen for subsequent biochemical analysis. A cross-section of the LV at the level of the papillary muscles was fixed in 4% PFA in PBS overnight at 4°C followed by preservation in a 30% sucrose solution and embedding in OCT.

### Histological Analyses

Frozen tissue sections were cut at a thickness of 10 μm. To evaluate total ventricular fibrotic area, cardiac sections were stained with Picrosirius red dye (0.1% Direct Red 80 in saturated picric acid solution) for 1 hour at room temperature. Whole-heart images were acquired using the image stitching feature on a Keyence BZ-X710 All-in-One Fluorescence Microscope. Fibrotic area was determined by quantifying the ratio of positively stained (red) pixels to the total pixel number of each section (reported as percent collagen content) using ImageJ. In addition, individual cardiomyocyte hypertrophy was assessed by staining cardiac sections with Alexa Fluor 647 wheat germ agglutinin (ThermoFisher W32466) according to manufacturer instructions. Briefly, sections were incubated in a 5.0 μg/mL WGA conjugate solution in HBSS for 10 minutes at room temperature, followed by mounting in ProLong™ Glass Antifade Mountant (ThermoFisher P36980). Images were acquired at 40x magnification with the left ventricular free wall using the Keyence BZ-X710 microscope, and individual cardiomyocyte area was quantified using ImageJ.

### Patient outcomes associated with drug treatments

To examine whether drugs were correlated with a reduction in cardiac adverse events associated with hypertrophy, we analyzed data from two separate clinical databases. As the first patient cohort, data was collected on November 30^th^, 2023 from the FDA’s Adverse Event Reporting System using the AERSMine tool^32^. The Adverse Event Reporting System is a multi-cohort database containing 20 million reports of adverse events from healthcare providers and consumers across the United States. As a second patient cohort, on January 2^nd^, 2024 we collected data from the University of Virginia Health System Network using the TriNetX tool, which provided access to electronic medical records from approximately 1.7 million patients.

This retrospective study is exempt from informed consent. The data reviewed is a secondary analysis of existing data, does not involve intervention or interaction with human subjects, and is de-identified per the de-identification standard defined in Section §164.514(a) of the HIPAA Privacy Rule. The process by which the data is de-identified is attested to through a formal determination by a qualified expert as defined in Section §164.514(b)(1) of the HIPAA Privacy Rule. This formal determination by a qualified expert refreshed on December 2020.

### Statistics

All data are presented as the mean ± SEM. Analysis of experimental conditions considered two distinct cardiomyocyte isolations and multiple conditions, with multiple wells in each experiment. For this reason, a two-way ANOVA followed by Dunnett’s test for multiple comparisons was selected for statistical calculation. In vivo data were analyzed by ordinary one-way ANOVA followed by Tukey’s multiple comparisons test. GraphPad Prism 9 was used for statistical analysis. For all analyses, a p-value <0.05 was considered statistically significant.

## RESULTS

### Identified drug pathways driving cardiomyocyte hypertrophy

Previously, we performed a microscopy-based screen for drugs that decreased hypertrophy of cardiomyocytes in the presence of phenylephrine^12^. In that study, 94 of 3241 small molecule compounds decreased both cell area and ANP protein expression by at least 70% and did not exhibit cardiotoxicity. In a previous study, we identified that the hypertrophy signaling model was directly targeted by 32 of these 94 drugs, and that the model could mechanistically predict reduction in hypertrophy with 88% (28 of 32) prediction accuracy ^28^. The remaining 62 of 94 compounds do not target this previous hypertrophy model and presumably inhibit cardiomyocyte hypertrophy through less-characterized mechanisms. Therefore, in this study we sought to identify novel pathways linking the remaining 62 compounds to hypertrophy. To identify new drug pathways, we developed a logic-based network predictor of drug pathways (LogiRx) that identifies and simulates molecular pathways that mediate how drugs regulate cell phenotype.

As a first application, we employed LogiRx to identify proteins that are directly targeted by the 62 anti-hypertrophic compounds from our prior screen^12^. LogiRx analysis mapped 28 of the 62 compounds to 79 protein targets in the OmniPath protein interaction database (**Figure 1A**). 11 of 28 compounds were successfully mapped to the hypertrophy network via 214 candidate drug pathways (**Figure 1B**). These pathways link 11 drugs and their drug targets to 89 proteins in the network, covering 42% of the network. Most of the candidate drug pathways (76%) connect the drug target to the network via an intermediate protein that is not in the original network model, demonstrating the value of pathway inference. Together, the 214 candidate drug-target-network pathways predict mechanisms that may help explain the experimentally-demonstrated anti-hypertrophic effect of these 11 compounds.

### Logic-based network expansion to predict how drugs regulate hypertrophy

To predict the impact of these drug pathways on cardiomyocyte hypertrophy, logic-based differential equation model variants were created from LogiRx that each simulate the effect of incorporating individual pathway (**Figure 1C**). Each model variant was tested by simulating inhibition of the drug target in the presence of hypertrophic agonist phenylephrine. Most drug pathways were predicted to strongly inhibit hypertrophy (**Figure 2A**). However, some drug pathways reduced the model accuracy when tested against a previous set of 450 experiments, indicating that they are not fully compatible with the mechanisms established in the validated network model (**Figure 2A**).

**Figure 2.**
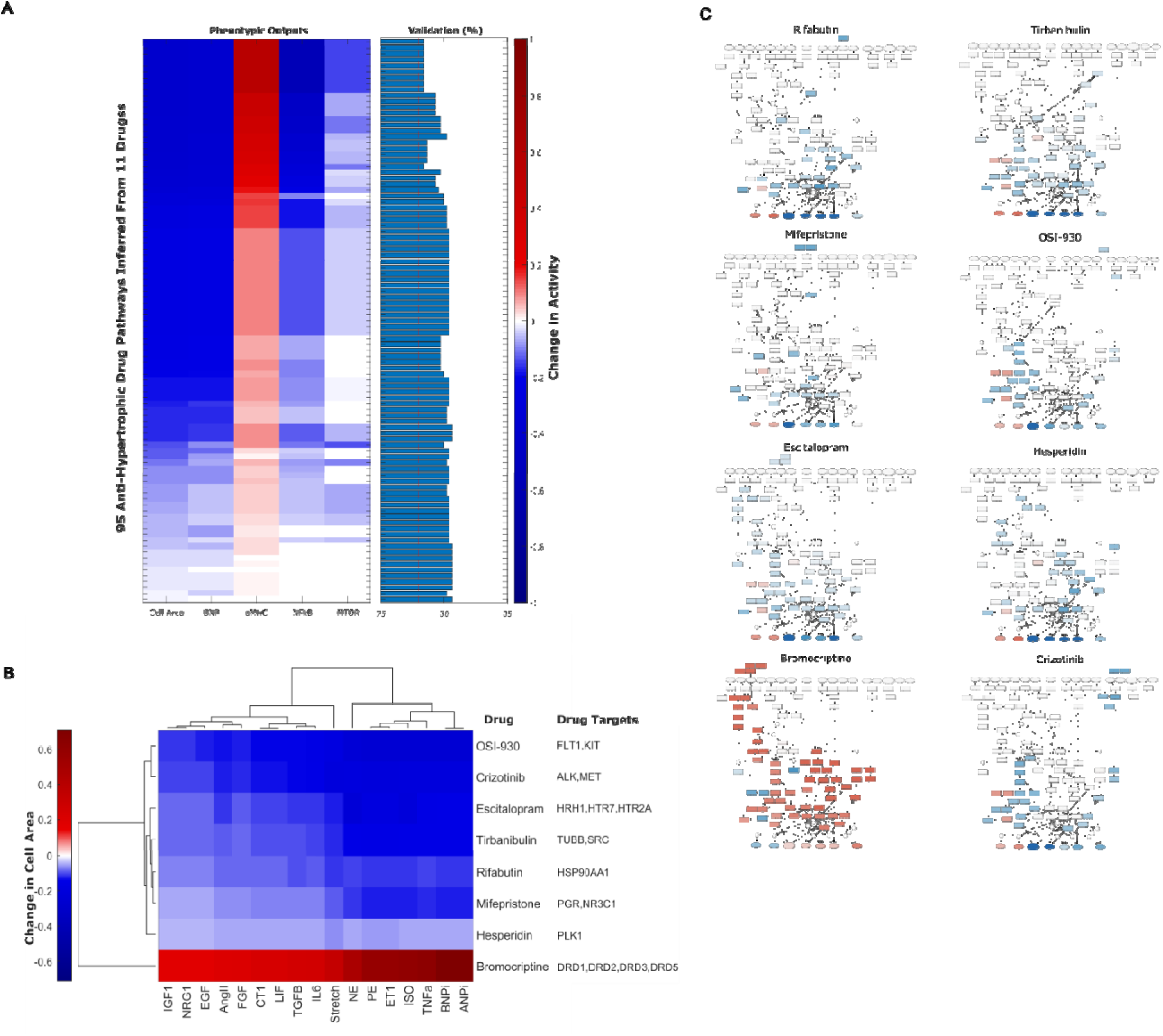
LogiRx-based network model expansion predicts pathways by which drugs affect cardiomyocyte hypertrophy. A) Hypertrophy network variants expanded by individual drug pathways were simulated for their effect on five hypertrophic outputs, as well as validated against 450 experiments from prior literature. Drug pathway annotation is provided in **Supplementary Figure S1**. B) 8 drugs were predicted to affect cardiomyocyte hypertrophy with varying efficacy across 17 biochemical environments. C) Drugs were predicted to inhibit hypertrophy via distinct regional modulation of the hypertrophy signaling network, indicating distinct inhibitory mechanisms.

Drug pathways corresponding to 8 of the 11 drugs were predicted to inhibit phenylephrine-induced hypertrophy while maintaining model accuracy above 78%. New LogiRx-expanded drug models were developed combining all relevant validated pathways for each of the 8 drugs. The hypertrophic effect of each drug was simulated using a pharmacological inhibition model based on known drug-target binding mechanisms^27^. Seven of these eight drugs were predicted to be anti-hypertrophic, with context-dependent efficacy depending on the hypertrophic stimulus (**Figure 2B**). 7 of 8 drugs regulated hypertrophy via multiple protein targets.

Seven drugs were predicted by the LogiRx-expanded network model to inhibit hypertrophy – OSI-930, crizotinib, escitalopram, tirbanibulin, rifabutin, mifepristone, and hesperidin, representing a variety of drug targets and clinical applications. OSI-930 is an FLT1 and KIT inhibitor that is designed to target cancer cell proliferation in solid tumors^33^. Crizotinib, an ALK and MET inhibitor, is used to treat non-small cell lung cancer but has known cardiac toxicity^34,35^. Escitalopram targets SERT and is used to treat depression and anxiety^36,37^. Tirbanibulin targets TUBB and SRC to treat actinic keratosis, and is being investigated as a potential treatment for acute myeloid leukemia^38^. Rifabutin inhibits HSP90AA1 and is an antibiotic preventing mycobacterium avium complex in HIV patients^39^. Mifepristone, which targets both PGR and NR3C1, is commonly used as an abortifacient, but is additionally used to treat diabetic patients with Cushing’s syndrome^40^. Hesperidin is a flavonoid and natural supplement currently investigated for anti-inflammatory benefits^41^. Similar to their target diversity, these 8 drugs were predicted to inhibit cardiomyocyte hypertrophy through distinct areas of the hypertrophy network (**Figure 2C**).

### Experiments in cardiomyocytes validate that escitalopram and mifepristone inhibit hypertrophy

We further prioritized escitalopram, tirbanibulin, mifepristone, OSI-930, and rifabutin for experimental validation based on their relative cardiac safety^33,42–47^ and approval status. Neonatal rat cardiomyocytes were treated with PE along with each of these five drugs for 48 h. Consistent with the LogiRx model predictions, tirbanibulin, escitalopram, and mifepristone inhibited PE-induced cardiomyocyte hypertrophy (**Figure 3A-B**). In contrast, OSI-930 and rifabutin did not inhibit hypertrophy even at increased concentrations.

**Figure 3.**
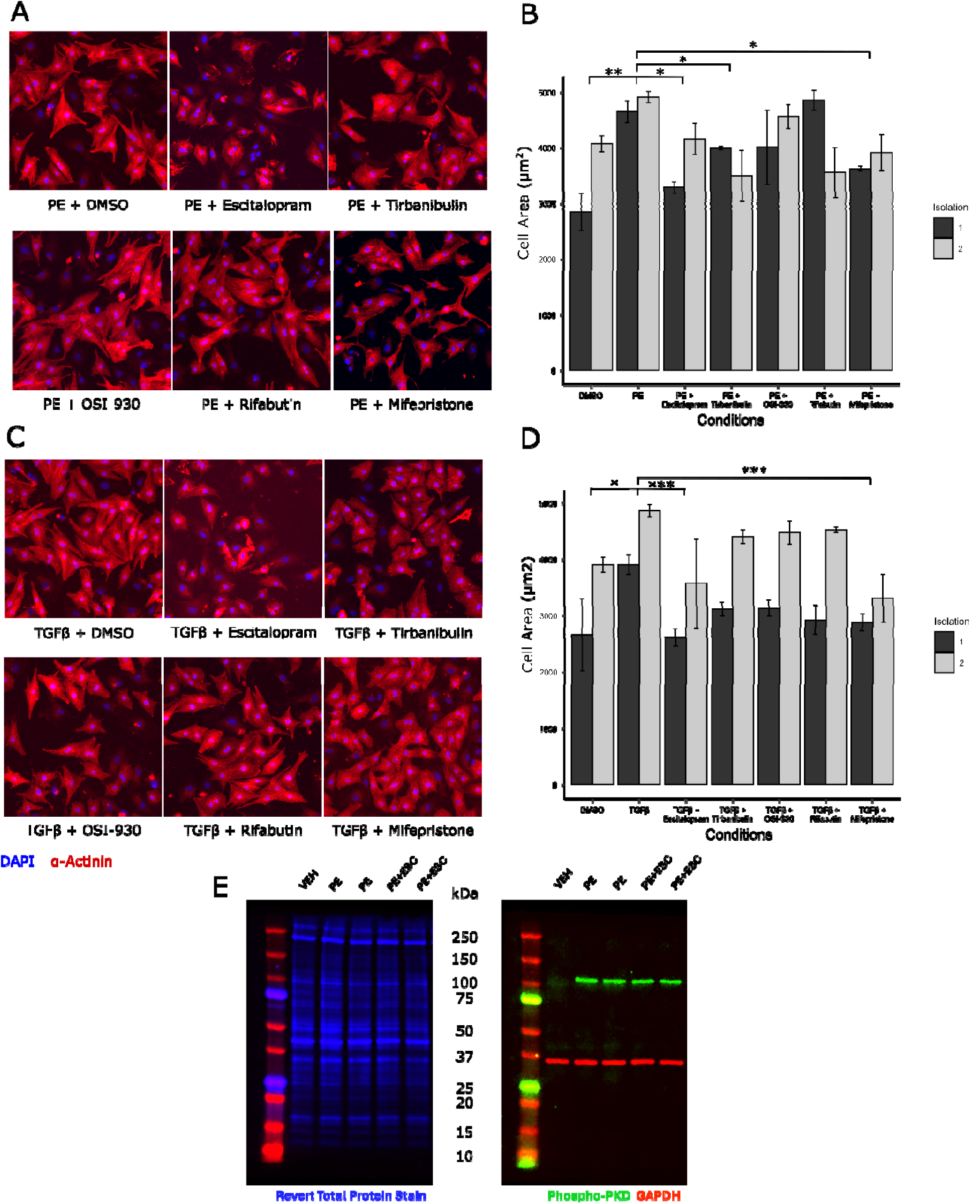
Escitalopram and mifepristone inhibit cardiomyocyte hypertrophy induced by either phenylephrine or TGFβ. The effect of five drugs on PE-induced hypertrophy of neonatal rat cardiomyocytes is shown in A) representative images or B) quantification of automated segmentation. Escitalopram (20 µM), mifepristone (6 µM), and tirbanibulin (6 µM) significantly prevented PE-induced cell area increase over a 48-hour period following treatment. The effect of these five drugs on TGFβ-induced hypertrophy is shown as C) representative images or D) quantification of cardiomyocyte cell area by automated segmentation. Escitalopram (20 µM) and mifepristone (6 µM) significantly prevented TGFβ-induced increase in cell area. E) Western blot of total protein and phospho-PKD in response to PE and/or escitalopram. Two cell isolations were used for each experiment, and a two-way ANOVA was performed followed by a Dunnett’s post-hoc test. Error bar indicates SEM, * p<0.05, ** p<0.01, *** p<0.001.

While the network analysis in **Figure 2C** predicted that escitalopram would work via downstream hypertrophic pathways that are relatively context-independent, an alternative hypothesis may suggest that escitalopram inhibits alpha-1 adrenergic receptor (α1AR) directly in response to phenylephrine. We tested this hypothesis by introducing an alternative hypertrophic stimulus TGFβ, as well as directly measuring phosphorylation of protein kinase D (PKD), which is proximal to the α1 adrenergic receptor in cardiomyocytes. TGFβ-induced hypertrophy was inhibited by both escitalopram and mifepristone (**Figure 3C-D**). Further, while we observed a robust increase in phospho-PKD levels in cardiomyocytes treated with phenylephrine stimulation, no reduction in PKD phosphorylation was observed in cardiomyocytes co-treated with escitalopram (**Figure 3E**). Together, these data indicate that escitalopram antagonizes hypertrophy through downstream mechanisms distinct from blockade of the α1AR.

To identify how mifepristone inhibits cardiomyocyte hypertrophy, we performed mechanistic subnetwork analysis. Although the LogiRx network model included drug pathways for mifepristone via both progesterone receptors and glucocorticoid receptors, LogiRx predicted that mifepristone inhibits cardiomyocyte hypertrophy predominantly via inhibition of the glucocorticoid receptor via LCK/ERK and CEBPβ (**Supplementary Figure S2A-B**). We experimentally validated that the glucocorticoid receptor antagonist AL082D06 phenocopied the effects of mifepristone by preventing hypertrophy with PHE or TGFβ (**Supplementary Figure S2C-D**).

### Escitalopram inhibits hypertrophy through serotonin receptor and not transporter

As the last stage of LogiRx, we performed mechanistic subnetwork analysis to identify pathways that mediate escitalopram inhibition of cardiomyocyte hypertrophy. We simulated the network-wide response to escitalopram, followed by simulations of network-wide knockdown of each network node in the presence and absence of escitalopram. The intersection of these two analyses resulted in a mechanistic subnetwork that predicted how escitalopram inhibits cardiomyocyte hypertrophy (**Figure 4A**). This subnetwork predicted that escitalopram inhibits hypertrophy through the serotonin receptor HT2RA, a non-canonical target of escitalopram, rather than as a selective serotonin reuptake inhibitor (SSRI) via the transporter SERT (or SLC6A4). Further, the LogiRx model predicted that the escitalopram-HTR2A pathway acts by inhibiting G-protein signaling to PI3Kγ (**Figure 4A-B**). The model also correctly predicted how αMHC and βMHC gene expression respond to escitalopram (**Supplementary Figure S3**).

**Figure 4.**
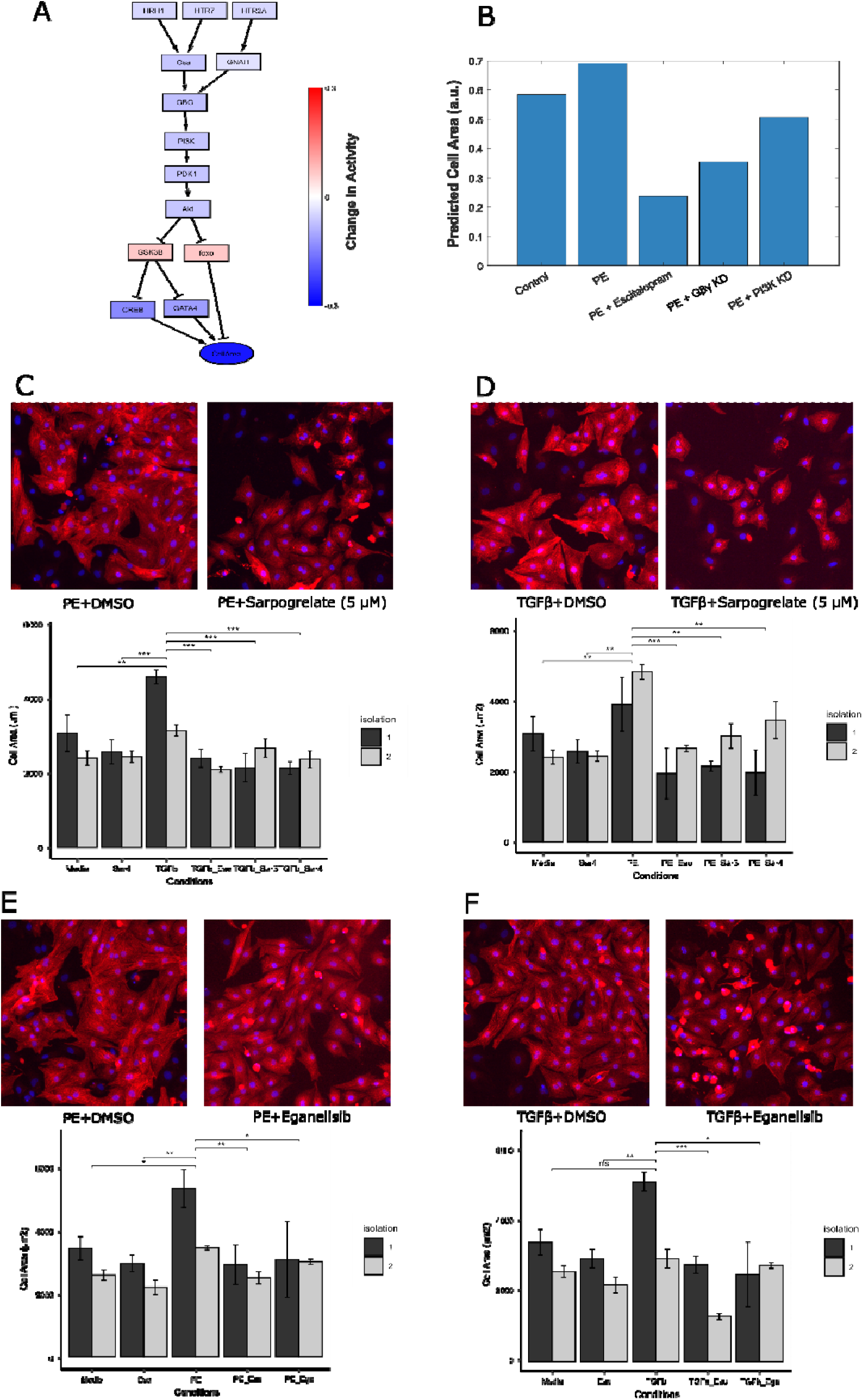
Escitalopram prevents cardiomyocyte hypertrophy by inhibiting serotonin receptors. A) Mechanistic subnetwork analysis to predict how escitalopram suppresses cardiomyocyte hypertrophy. B) Simulations of PE-induced hypertrophy and modulation by escitalopram, Gβγ inhibition, or PI3Kγ inhibition. Serotonin receptor inhibitor sarpogrelate hydrochloride (5 µM) significantly prevents C) PE- and D) TGFβ-induced cardiomyocyte hypertrophy 48 hours post treatment, as shown by representative images and quantification of automated segmentation. PI3Kγ inhibitor eganelisib (20 µM) significantly prevents E) PE- and F) TGFβ-induced cardiomyocyte hypertrophy 48 hours post treatment, as shown by representative images and quantification of automated segmentation. Two cell isolations were used for each experiment, and a two-way ANOVA was performed followed by a Dunnett’s post-hoc test. Error bar indicates SEM, * p<0.05, ** p<0.01, *** p<0.001.

To experimentally validate these LogiRx predictions, we used sarpogrelate hydrochloride which selectively inhibits HTR2A but not SERT. Prior literature^48,49^ demonstrates that sarpogrelate inhibits PE-induced cardiomyocyte hypertrophy both in vitro and in vivo. Our studies confirm that sarpogrelate prevents both PE- and further demonstrates its efficacy against TGFβ-induced cardiomyocyte hypertrophy (**Figure 4C-D**). Downstream of HTR2A, our mechanistic subnetwork predicted Gβγ and PI3K to mediate anti-hypertrophic escitalopram activity. Gβγ directly binds and activates PI3Kγ^50^. To experimentally validate this prediction, we selectively inhibited PI3Kγ with the antitumor drug eganelisib. Eganelisib prevented both PE- and TGFβ-induced cardiomyocyte hypertrophy (**Figure 4E-F**), further validating the non-canonical escitalopram-HTR2A-PI3Kγ pathway predicted by LogiRx.

### Escitalopram inhibits cardiomyocyte hypertrophy *in vivo*

To test whether escitalopram also reduces cardiomyocyte hypertrophy in vivo, escitalopram was delivered to mice subjected to the angiotensin II and phenylephrine (AngII/PE) as a mouse model of cardiac hypertrophy and fibrosis. Angiotensin II (1.5 mg/kg/day) and phenylephrine (50 mg/kg/day) were delivered continuously for 14 days through an osmotic minipump as previously described^51^; daily escitalopram was administered via intraperitoneal injection at 10 mg/kg/day beginning the day after pump implantation (**Figure 5A**). Mice subjected to AngII/PE for 14 days exhibited significant global cardiac hypertrophy, as indicated by heart weight to body weight ratio (HW:BW), as well as a significant increase in LV mass in particular (LV:TL) (**Figure 5B-C**). Furthermore, AngII/PE challenge induced dramatic fibrosis in the LV free wall, which likely also contributed to the increase in overall LV mass (**Figure 5D-E**). Cardiac structure and systolic and diastolic function were evaluated by echocardiography, revealing an increase in LV wall thickness in diastole, which is consistent with the increased HW:BW ratio (**Figure 5F**). Furthermore, AngII/PE mice demonstrated trends towards increases in systolic function, measured by ejection fraction, as well as increases in the E/E’ ratio, which is indicative of impaired LV relaxation or diastolic dysfunction (**Figure 5G-H**).

**Figure 5.**
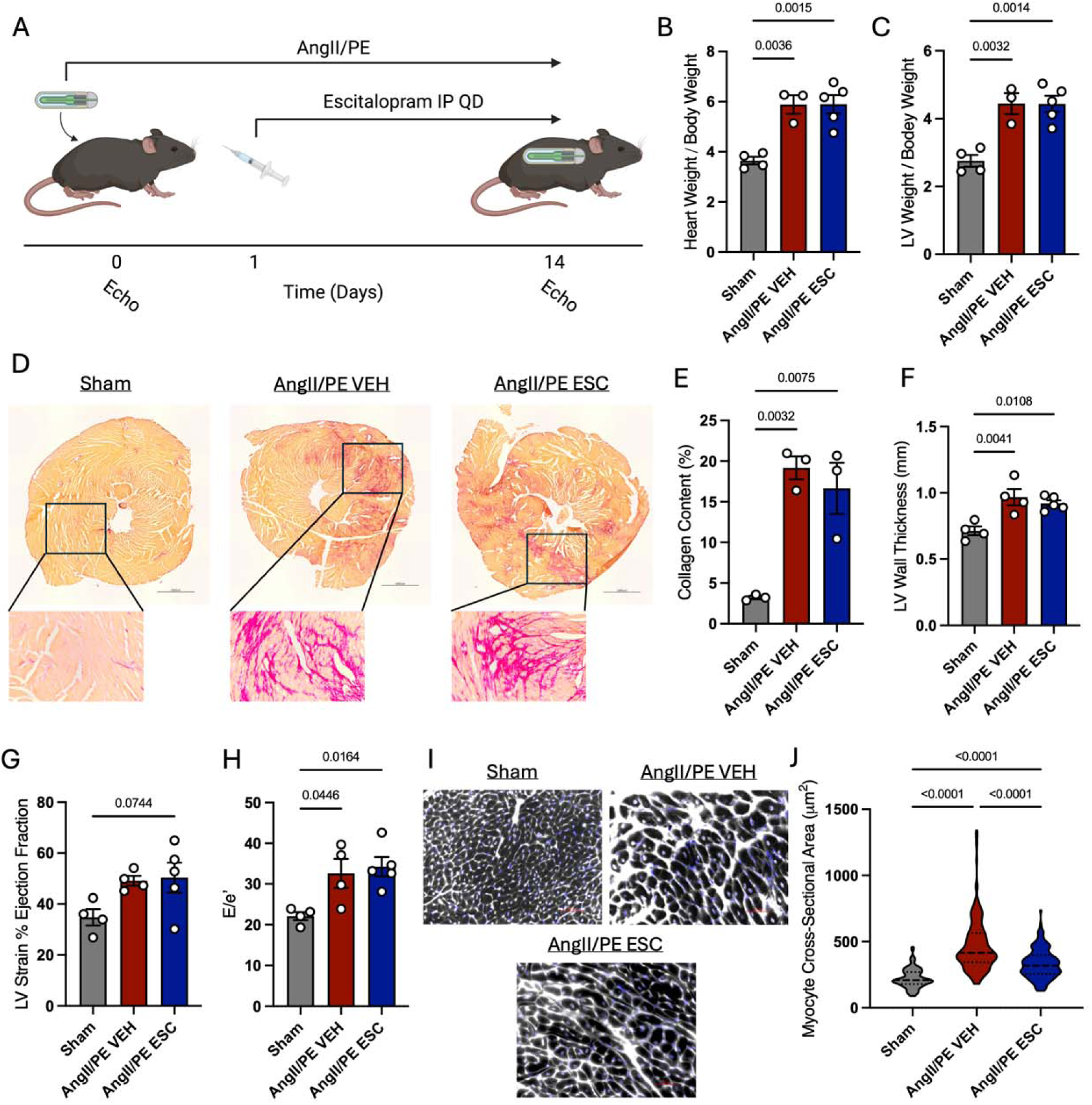
Escitalopram Attenuates Cardiomyocyte Hypertrophy in AngII/PE Mice. A. Schematic of cardiac injury model and escitalopram administration. Mice were continuously delivered AngII/PE through an osmotic minipump to induce cardiac injury and fibrosis, and injected with 10 mg/kg/day escitalopram or vehicle control for 14 days. Quantification of HW:BW (B) and LV:BW (C) ratios. D. Representative picrosirius red staining of heart cross-sectional tissues in sham mice and AngII/PE mice treated with vehicle or escitalopram, and quantification of percent collagen content (E). Results of echocardiographic analyses of AngII/PE mice, showing the left ventricular wall thickness in diastole (F), percent ejection fraction derived from strain analysis (G), and the ratio of early mitral valve flow velocity to mitral annulus velocity by tissue Doppler (E/e’) indicating diastolic function. I. Wheat germ agglutinin staining of tissue sections, with quantification of cardiomyocyte cross-sectional area (J). Statistical analyses performed by one-way ANOVA with Tukey’s post hoc multiple comparisons test.

Some mice treated with AngII/PE also received daily administration of escitalopram delivered via intraperitoneal injection at 10 mg/kg/day. These animals showed no change in overall collagen content in the LV, suggesting a lack of an effect on the cardiac fibroblast population (**Figure 5D-E**). There was also no significant effect of escitalopram on global cardiac hypertrophy or any functional parameters assessed by echocardiography (**Figure 5B-C,F-H**). However, imaging of cardiomyocyte sections stained with WGA revealed a modest but significant reduction in individual cardiomyocyte hypertrophy in response to treatment with escitalopram (**Figure 5I-J**). In summary, while escitalopram did not offer significant protection against global cardiac hypertrophy or dysfunction in this severe injury model, escitalopram attenuated cardiomyocyte hypertrophy in vivo, consistent with the antihypertrophic effect of escitalopram predicted by LogiRx and measured in cultured cardiomyocytes.

### Cardiac outcomes for patients prescribed escitalopram

Based on consistent anti-hypertrophic roles of escitalopram shown by LogiRx, cultured rat cardiomyocytes, and mouse cardiomyocytes in vivo, we asked whether patients prescribed escitalopram may also exhibit a reduced incidence of cardiac hypertrophy. We first mined patient reports from the FDA Adverse Event Reporting System for incidence of cardiac hypertrophy or failure among patients prescribed escitalopram or related SSRIs that are not known to target HTR2A. Among those diagnosed with depression, patients prescribed escitalopram (which inhibits both SERT and HTR2A) had a lower incidence of left ventricular or ventricular hypertrophy when compared to treatment with other SSRIs sertraline, venlafaxine, and paroxetine that inhibit SERT but not HTR2A (**Figure 6A**). An exception is patients treated with fluoxetine, who had lower incidence of cardiac failure compared to escitalopram treatment.

**Figure 6.**
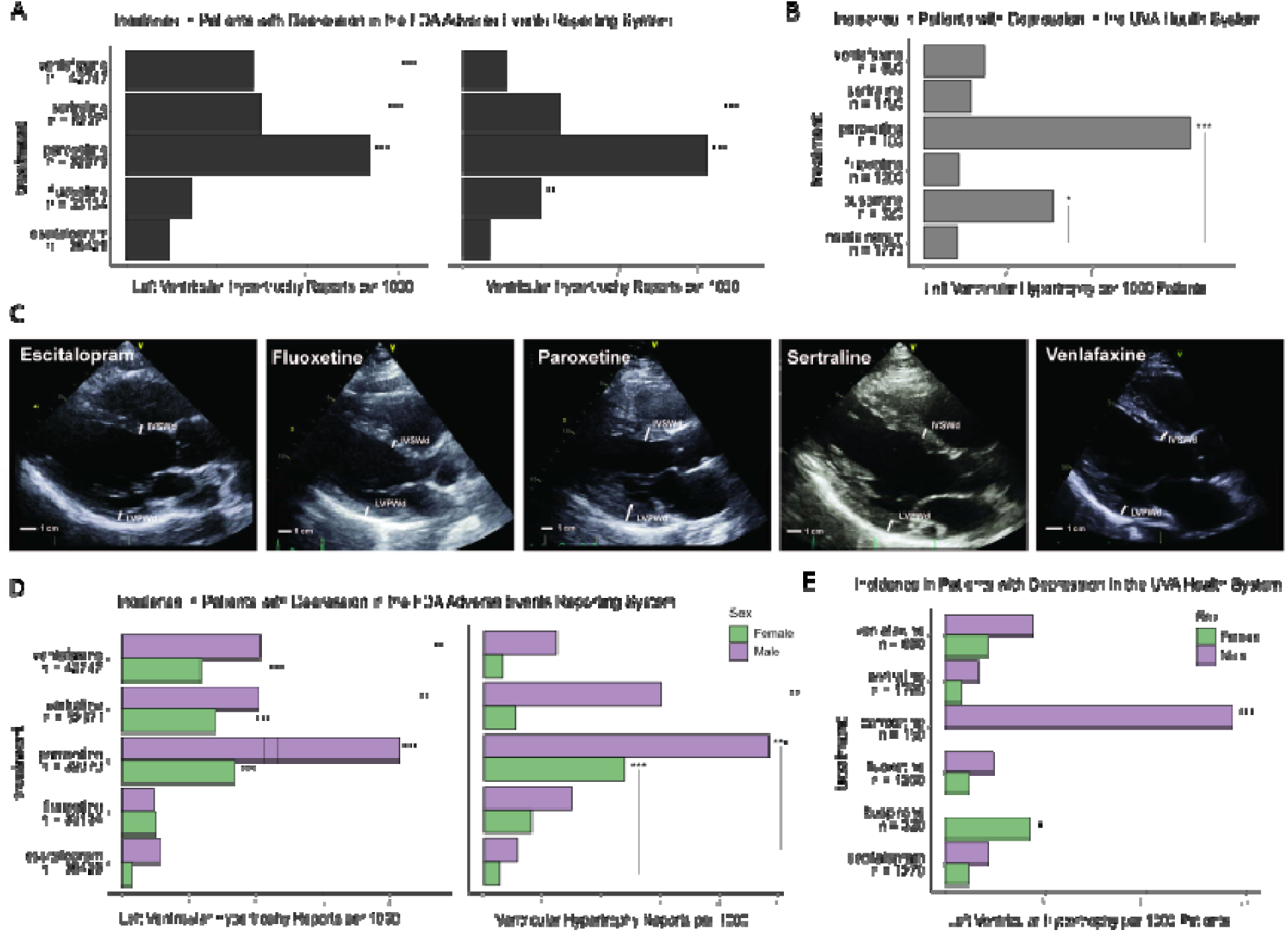
In two patient cohorts with depression, those prescribed escitalopram exhibit a lower incidence of cardiac hypertrophy than those prescribed other SSRIs. A) Reports of left ventricular hypertrophy (left) or ventricular hypertrophy (right) per 1000 patients are shown for patients with depression taking either escitalopram or another SSRI (fluoxetine, paroxetine, sertraline, or venlafaxine) in the FDA Adverse Events Reporting System. B) Diagnoses of left ventricular hypertrophy per 1000 patients are shown for patients with depression taking either escitalopram or another SSRI in the U. Virginia Health System. C) Verification of echocardiograms from patients in the UVA Health System with hypertrophy (prescribed paroxetine) or without hypertrophy (prescribed escitalopram, fluoxetine, sertraline, or venlafaxine). D) FDA adverse event reports of left ventricular hypertrophy (left) or ventricular hypertrophy (right) per 1000 patients are further divided into incidence by sex. E) Diagnoses of left ventricular hypertrophy in the UVA Health System, divided into incidence by sex. In the UVA Health System, no female patients prescribed paroxetine and no male patients prescribed busipirone exhibited left ventricular hypertrophy. * p<0.05, ** p<0.01, *** p<0.001.

To validate these findings in an independent patient population with more structured data collection, we mined electronic health records from the University of Virginia (UVA) Health System. Among UVA patients diagnosed with depression, those prescribed escitalopram exhibited a statistically lower incidence of left ventricular hypertrophy than those prescribed SSRI paroxetine as well as the serotonin receptor agonist buspirone (**Figure 6B**). To verify the hypertrophy annotation from this analysis, we examined echocardiograms from de-identified patients using the same drug and diagnosis queries. These confirm qualitatively larger interventricular septum thickness (IVSD) and left ventricular posterior wall end diastole (LVPWD) in a patient prescribed paroxetene and diagnosed with hypertrophy than echocardiograms from patients prescribed escitalopram or other SSRIs not diagnosed with hypertrophy (**Figure 6C**). Stratifying patients in the FDA (**Figure 6D**) and UVA (**Figure 6E**) databases by sex, both male and female patients with depression prescribed escitalopram exhibit less hypertrophy than that seen with two or more SSRIs. A reduced rate of hypertrophy was also seen in patients in the FDA database with anxiety on escitalopram compared with other SSRIs (**Supplementary Figure S4**).

## Discussion

Despite a strong association between heart failure and hypertrophy, drugs that directly treat hypertrophy are limited^52^. As seen with sildenafil, drug repurposing may provide a rapid translational avenue to identify new therapeutic strategies for cardiac hypertrophy and heart failure^53,54^. Drug repurposing requires sophisticated screening for efficacy but also methods to identify how effective drugs work. In this study we developed a new mechanistic machine learning method, LogiRx, that predicts new pathways that map from drugs to a mechanistic logic-based model of a signaling network. We applied LogiRx to hits from our previous experimental compound screen^12^ and predicted the pathways by which 7 of these drugs modulate cardiomyocyte hypertrophy. Experiments in cultured cardiomyocytes validated the context-independent antihypertrophic role of two drugs, escitalopram and mifepristone predicted by LogiRx. Pathways that mediate the effects of escitalopram and mifepristone were predicted by subnetwork analysis, and these predictions were further experimentally validated (escitalopram: serotonin receptors and Gβγ; mifepristone: glucocorticoid receptor). Analysis of electronic health records validate these in silico and in vitro results findings, demonstrating that escitalopram is associated with a lower incidence of cardiac hypertrophy in depression or anxiety patients compared to multiple SSRIs in two large independent populations. Together, these results suggest potential repurposing avenues for escitalopram and mifepristone in cardiac hypertrophy.

LogiRx identifies regulatory signaling pathways and dynamic mechanistic predictions that map from drugs to an established signaling network. Several other computational methods construct signaling pathways from both protein-interaction and/or gene expression data^55–57^. Notably, CARNIVAL combines gene expression with protein interaction data to predict upstream signaling pathways that drive changes gene expression changes^58^. LogiRx provides additional benefits of its logic-based differential equation framework that is highly validated for cardiomyocyte hypertrophy^19,20,28,59^ to provide dynamic, mechanistic simulations of perturbation experiments (**Figure 2, 4**). The performance of LogiRx depends on the accuracy and completeness of drug, target, and network model resources, as well as the characteristics of the validation experiments. These limitations and potential improvements are described in **Supplementary Table S1**. LogiRx may be most readily applied to settings where annotated drugs have been used to screen cellular phenotype or gene expression, and there is also a validated mechanistic network model that predicts regulation of that cell response. For example, LogiRx may be amenable for integrating drug screens and mechanistic network models in areas such as myofibroblast transformation for fibrosis^60,61^ and proliferation for breast cancer^62,63^. Challenges to this approach arise when there is insufficient characterization of drug targets, inadequate relevance of the drug targets to the network model, or an inadequately validated mechanistic network model.

Escitalopram (Lexapro) is a commonly prescribed SSRI (>30M prescriptions in the US) with a well-known safety profile^36,37^. Less well understood is the role of SSRIs in cardiac disease. Escitalopram was shown to prevent a diabetic model of cardiac hypertrophy, however the mechanisms remain poorly understood^64^. Our LogiRx model predicts that escitalopram inhibits cardiomyocyte hypertrophy through its off-target HTR2A. This predicted mechanism is supported by our understanding of the role of serotonin receptors HTR2A and HTR2B^65–67^ in cardiomyocyte hypertrophy. Escitalopram decreased cardiomyocyte size in cell culture, mice, and was associated with lower incidence of ventricular hypertrophy in patients, but escitalopram did not reduce global heart weight in the mice. We hypothesize that this difference may be due marked fibrosis in this mouse model and partial attenuation of cardiomyocyte hypertrophy by escitalopram (**Figure 5D**).

Paroxetine, another SSRI that more specifically targets SERT, was previously shown to prevent left ventricular dysfunction following myocardial infarction through an off-target GRK2 mechanism^68^. GRK2 is unlikely to mediate the increased incidence of hypertrophy we found associated with patients prescribed paroxetine, because the clinical concentration is below the IC50 for GRK2^69,70^. Escitalopram off-target of HT2A and paroxetine off-target of GRK2 indicates that other SSRIs may have off-targets that contribute to the differential association with cardiac hypertrophy and failure we observed in patients with depression or anxiety. Further, paroxetine derivatives with enhanced affinity for GRK2 demonstrate the potential for selective repurposing towards “off targets”^71,72^.

While mifepristone is best known as a progesterone inhibitor, it was originally discovered in a search for glucocorticoid receptor antagonists and is used in Cushing’s syndrome with hyperglycemia^40^. Glucocorticoids and their receptors have a complex role in cardiovascular health. Glucocorticoid stimulation induces hypertrophy in primary rat cardiomyocytes^73,74^, yet mice with cardiomyocyte-specific deletion of glucocorticoid receptor have increased cardiac hypertrophy^75^ and direct glucocorticoid receptor inhibition can enhance cardiomyocyte proliferation^76^. LogiRx predicted that mifepristone inhibits cardiomyocyte hypertrophy through a glucocorticoid receptor - LCK – ERK pathway, which we validated in two contexts in cultured cardiomyocytes. The implications of mifepristone for repurposing against cardiac hypertrophy warrant further investigation.

Machine learning methods that not only predict efficacy but also pathway mechanisms are greatly needed. Here, we developed a mechanistic machine learning method LogiRx to discover pathways by which escitalopram and mifepristone inhibit cardiomyocyte hypertrophy. These pathways were validated in cultured cells in two contexts. Escitalopram decreased cardiomyocyte hypertrophy in vivo and was associated with a lower incidence of cardiac hypertrophy in two patient cohorts. Together, these studies demonstrate the ability of LogiRx to predict drug efficacy and mechanisms that guide new experiments.

## Acknowledgments

We thank Mohammad Fallahi-Sichani for use of his Perkin Elmer Operetta, statistical consulting from Marieke Jones, manuscript feedback from Kevin Janes and Jason Papin, and Malcolm O’Malley for naming LogiRx.

## References

1. Frey, N., Katus, H. A., Olson, E. N. & Hill, J. A. Hypertrophy of the Heart. Circulation 109, 1580–1589 (2004).

2. Samak, M. et al. Cardiac Hypertrophy: An Introduction to Molecular and Cellular Basis. Med. Sci. Monit. Basic Res. 22, 75–79 (2016).

3. Nomura, S. et al. Cardiomyocyte gene programs encoding morphological and functional signatures in cardiac hypertrophy and failure. Nat. Commun. 9, 4435 (2018).

4. Marian, A. J. 23 - Heart Failure as a Consequence of Hypertrophic Cardiomyopathy. in Heart Failure: a Companion to Braunwald’s Heart Disease (Fourth Edition) (eds. Felker, G. M. & Mann, D. L.) 311–321.e6 (Elsevier, Philadelphia, 2020). doi:10.1016/B978-0-323-60987-6.00023-5.

5. Bazgir, F. et al. The Microenvironment of the Pathogenesis of Cardiac Hypertrophy. Cells 12, 1780 (2023).

6. Xie, M., Burchfield, J. S. & Hill, J. A. Pathological Ventricular Remodeling: Mechanisms: Part 1 of 2. Circulation 128, 388–400 (2013).

7. Tham, Y. K., Bernardo, B. C., Ooi, J. Y. Y., Weeks, K. L. & McMullen, J. R. Pathophysiology of cardiac hypertrophy and heart failure: signaling pathways and novel therapeutic targets. Arch. Toxicol. 89, 1401–1438 (2015).

8. Tsao, C. W. et al. Heart Disease and Stroke Statistics-2023 Update: A Report From the American Heart Association. Circulation 147, e93–e621 (2023).

9. Schubert, M., Hansen, S., Leefmann, J. & Guan, K. Repurposing Antidiabetic Drugs for Cardiovascular Disease. Front. Physiol. 11, 568632 (2020).

10. Onódi, Z., Koch, S., Rubinstein, J., Ferdinandy, P. & Varga, Z. V. Drug repurposing for cardiovascular diseases: New targets and indications for probenecid. Br. J. Pharmacol. 180, 685–700 (2023).

11. Xie, Y. et al. Mechanisms of SGLT2 Inhibitors in Heart Failure and Their Clinical Value. J. Cardiovasc. Pharmacol. 81, 4–14 (2023).

12. Reid, B. G. et al. Discovery of novel small molecule inhibitors of cardiac hypertrophy using high throughput, high content imaging. J. Mol. Cell. Cardiol. 97, 106–113 (2016).

13. Dukinfield, M. et al. Repurposing an anti-cancer agent for the treatment of hypertrophic heart disease. J. Pathol. 249, 523–535 (2019).

14. Escudero, D. S. et al. PDE5 inhibition improves cardiac morphology and function in SHR by reducing NHE1 activity: Repurposing Sildenafil for the treatment of hypertensive cardiac hypertrophy. Eur. J. Pharmacol. 891, 173724 (2021).

15. Wang, Z. et al. A high-throughput drug screening identifies luteolin as a therapeutic candidate for pathological cardiac hypertrophy and heart failure. Front. Cardiovasc. Med. 10, 1130635 (2023).

16. Nakamura, M. & Sadoshima, J. Mechanisms of physiological and pathological cardiac hypertrophy. Nat. Rev. Cardiol. 15, 387–407 (2018).

17. Sun, J., Yang, J., Chi, J., Ding, X. & Lv, N. Identification of drug repurposing candidates based on a miRNA-mediated drug and pathway network for cardiac hypertrophy and acute myocardial infarction. Hum. Genomics 12, 52 (2018).

18. Rai, A., Kumar, V., Jerath, G., Kartha, C. C. & Ramakrishnan, V. Mapping drug-target interactions and synergy in multi-molecular therapeutics for pressure-overload cardiac hypertrophy. NPJ Syst. Biol. Appl. 7, 11 (2021).

19. Ryall, K. et al. Network reconstruction and systems analysis of cardiac myocyte hypertrophy signaling. J. Biol. Chem. 287, 42259–42268 (2012).

20. Khalilimeybodi, A., Paap, A. M., Christiansen, S. L. M. & Saucerman, J. J. Context-specific network modeling identifies new crosstalk in β-adrenergic cardiac hypertrophy. PLoS Comput. Biol. 16, (2020).

21. Wishart, D. S. et al. DrugBank 5.0: a major update to the DrugBank database for 2018. Nucleic Acids Res. 46, D1074–D1082 (2018).

22. Türei, D., Korcsmáros, T. & Saez-Rodriguez, J. OmniPath: guidelines and gateway for literature-curated signaling pathway resources. Nat. Methods 13, 966–967 (2016).

23. Ceccarelli, F., Turei, D., Gabor, A. & Saez-Rodriguez, J. Bringing data from curated pathway resources to Cytoscape with OmniPath. Bioinformatics 36, 2632–2633 (2020).

24. Gil, D. P., Law, J. N. & Murali, T. M. The PathLinker app: Connect the dots in protein interaction networks. F1000Research 6, 58 (2017).

25. Ritz, A. et al. Pathways on demand: automated reconstruction of human signaling networks. NPJ Syst. Biol. Appl. 2, 16002 (2016).

26. Shannon, P. et al. Cytoscape: A Software Environment for Integrated Models of Biomolecular Interaction Networks. Genome Res. 13, 2498–2504 (2003).

27. Zeigler, A. C. et al. Network model-based screen for FDA-approved drugs affecting cardiac fibrosis. CPT Pharmacomet. Syst. Pharmacol. 10, 377–388 (2021).

28. Eggertsen, T. G. & Saucerman, J. J. Virtual drug screen reveals context-dependent inhibition of cardiomyocyte hypertrophy. Br. J. Pharmacol. 180, 2721–2735 (2023).

29. Van de Graaf, M. W., Eggertsen, T. G., Zeigler, A. C., Tan, P. M. & Saucerman, J. J. Benchmarking of protein interaction databases for integration with manually reconstructed signalling network models. J. Physiol. (2023) doi:10.1113/JP284616.

30. Stirling, D. R. et al. CellProfiler 4: improvements in speed, utility and usability. BMC Bioinformatics 22, 433 (2021).

31. Bass, G. T. et al. Automated image analysis identifies signaling pathways regulating distinct signatures of cardiac myocyte hypertrophy. J. Mol. Cell. Cardiol. 52, 923–930 (2012).

32. Sarangdhar, M. et al. Data mining differential clinical outcomes associated with drug regimens using adverse event reporting data. Nat. Biotechnol. 34, 697–700 (2016).

33. Yap, T. A. et al. First-in-Human Phase I Trial of Two Schedules of OSI-930, a Novel Multikinase Inhibitor, Incorporating Translational Proof-of-Mechanism Studies. Clin. Cancer Res. Off. J. Am. Assoc. Cancer Res. 19, 10.1158/1078-0432.CCR-12–2258 (2013).

34. Chan, S. H. Y., Khatib, Y., Webley, S., Layton, D. & Salek, S. Identification of cardiotoxicity related to non-small cell lung cancer (NSCLC) treatments: A systematic review. Front. Pharmacol. 14, 1137983 (2023).

35. Xu, Z. et al. Inhibition of PRKAA/AMPK (Ser485/491) phosphorylation by crizotinib induces cardiotoxicity via perturbing autophagosome-lysosome fusion. Autophagy 20, 416–436.

36. Kasper, S. et al. Differences in the dynamics of serotonin reuptake transporter occupancy may explain superior clinical efficacy of escitalopram versus citalopram. Int. Clin. Psychopharmacol. 24, 119–125 (2009).

37. Sanchez, C., Reines, E. H. & Montgomery, S. A. A comparative review of escitalopram, paroxetine, and sertraline: are they all alike? Int. Clin. Psychopharmacol. 29, 185 (2014).

38. Kasner, M. T. et al. A phase Ib dose escalation study of oral monotherapy with KX2-391 in elderly patients with acute myeloid leukemia. Invest. New Drugs 40, 773–781 (2022).

39. Crabol, Y., Catherinot, E., Veziris, N., Jullien, V. & Lortholary, O. Rifabutin: where do we stand in 2016? J. Antimicrob. Chemother. 71, 1759–1771 (2016).

40. Pivonello, R., De Leo, M., Cozzolino, A. & Colao, A. The Treatment of Cushing’s Disease. Endocr. Rev. 36, 385–486 (2015).

41. Hajialyani, M. et al. Hesperidin as a Neuroprotective Agent: A Review of Animal and Clinical Evidence. Mol. Basel Switz. 24, 648 (2019).

42. Thase, M. E., Larsen, K. G., Reines, E. & Kennedy, S. H. The cardiovascular safety profile of escitalopram. Eur. Neuropsychopharmacol. J. Eur. Coll. Neuropsychopharmacol. 23, 1391–1400 (2013).

43. Kimura, K. et al. Cardiovascular adverse reactions associated with escitalopram in patients with underlying cardiovascular diseases: a systematic review and meta-analysis. Front. Psychiatry 14, 1248397 (2023).

44. Heppt, M. V. et al. Comparative Efficacy and Safety of Tirbanibulin for Actinic Keratosis of the Face and Scalp in Europe: A Systematic Review and Network Meta-Analysis of Randomized Controlled Trials. J. Clin. Med. 11, 1654 (2022).

45. Darpo, B., Bullingham, R., Combs, D. L., Ferber, G. & Hafez, K. Assessment of the cardiac safety and pharmacokinetics of a short course, twice daily dose of orally-administered mifepristone in healthy male subjects. Cardiol. J. 20, 152–160 (2013).

46. Mathew, S., Ticsa, M. S., Qadir, S., Rezene, A. & Khanna, D. Multiple Clinical Indications of Mifepristone: A Systematic Review. Cureus 15, e48372.

47. Phillips, M. C., Wald-Dickler, N., Loomis, K., Luna, B. M. & Spellberg, B. Pharmacology, Dosing, and Side Effects of Rifabutin as a Possible Therapy for Antibiotic-Resistant Acinetobacter Infections. Open Forum Infect. Dis. 7, ofaa460 (2020).

48. Ikeda, K. et al. The effects of sarpogrelate on cardiomyocyte hypertrophy. Life Sci. 67, 2991–2996 (2000).

49. Shimizu, K. et al. The Selective Serotonin 2A Receptor Antagonist Sarpogrelate Prevents Cardiac Hypertrophy and Systolic Dysfunction via Inhibition of the ERK1/2–GATA4 Signaling Pathway. Pharmaceuticals 14, 1268 (2021).

50. Vadas, O. et al. Molecular determinants of PI3Kγ-mediated activation downstream of G-protein– coupled receptors (GPCRs). Proc. Natl. Acad. Sci. 110, 18862–18867 (2013).

51. Aghajanian, H. et al. Targeting Cardiac Fibrosis with Engineered T cells. Nature 573, 430–433 (2019).

52. Palandri, C. et al. Pharmacological Management of Hypertrophic Cardiomyopathy: From Bench to Bedside. Drugs 82, 889–912 (2022).

53. Sapna, F. et al. Advancements in Heart Failure Management: A Comprehensive Narrative Review of Emerging Therapies. Cureus 15, e46486.

54. Abdelsayed, M., Kort, E. J., Jovinge, S. & Mercola, M. Repurposing drugs to treat cardiovascular disease in the era of precision medicine. Nat. Rev. Cardiol. 19, 751–764 (2022).

55. Kirouac, D. et al. Creating and analyzing pathway and protein interaction compendia for modelling signal transduction networks. BMC Syst. Biol. 6, 29 (2012).

56. Chen, B., Fan, W., Liu, J. & Wu, F.-X. Identifying protein complexes and functional modules--from static PPI networks to dynamic PPI networks. Brief. Bioinform. 15, 177–194 (2014).

57. Kabir, M. H., Patrick, R., Ho, J. W. K. & O’Connor, M. D. Identification of active signaling pathways by integrating gene expression and protein interaction data. BMC Syst. Biol. 12, 120 (2018).

58. Liu, A. et al. From expression footprints to causal pathways: contextualizing large signaling networks with CARNIVAL. NPJ Syst. Biol. Appl. 5, 40 (2019).

59. Frank, D. U., Sutcliffe, M. D. & Saucerman, J. J. Network-based predictions of in vivo cardiac hypertrophy. J. Mol. Cell. Cardiol. 121, 180–189 (2018).

60. Zeigler, A. C., Richardson, W. J., Holmes, J. W. & Saucerman, J. J. A computational model of cardiac fibroblast signaling predicts context-dependent drivers of myofibroblast differentiation. J. Mol. Cell. Cardiol. 94, 72–81 (2016).

61. Nelson, A. R., Christiansen, S. L., Naegle, K. M. & Saucerman, J. J. Logic-based mechanistic machine learning on high-content images reveals how drugs differentially regulate cardiac fibroblasts. Proc. Natl. Acad. Sci. U. S. A. 121, e2303513121 (2024).

62. Bouhaddou, M. et al. A mechanistic pan-cancer pathway model informed by multi-omics data interprets stochastic cell fate responses to drugs and mitogens. PLoS Comput. Biol. 14, e1005985 (2018).

63. Keenan, A. B. et al. The Library of Integrated Network-Based Cellular Signatures NIH Program: System-Level Cataloging of Human Cells Response to Perturbations. Cell Syst. 6, 13–24 (2018).

64. Ahmed, L. A., Shiha, N. A. & Attia, A. S. Escitalopram Ameliorates Cardiomyopathy in Type 2 Diabetic Rats via Modulation of Receptor for Advanced Glycation End Products and Its Downstream Signaling Cascades. Front. Pharmacol. 11, 579206 (2020).

65. Bush, E. et al. A small molecular activator of cardiac hypertrophy uncovered in a chemical screen for modifiers of the calcineurin signaling pathway. Proc. Natl. Acad. Sci. U. S. A. 101, 2870–2875 (2004).

66. Mialet-Perez, J. et al. Serotonin 5-HT2A receptor-mediated hypertrophy is negatively regulated by caveolin-3 in cardiomyoblasts and neonatal cardiomyocytes. J. Mol. Cell. Cardiol. 52, 502–510 (2012).

67. Gao, W. et al. HTR2A promotes the development of cardiac hypertrophy by activating PI3K-PDK1-AKT-mTOR signaling. Cell Stress Chaperones 25, 899–908 (2020).

68. Schumacher, S. M. et al. Paroxetine-mediated GRK2 inhibition reverses cardiac dysfunction and remodeling after myocardial infarction. Sci. Transl. Med. 7, 277ra31 (2015).

69. Thal, D. M. et al. Paroxetine is a direct inhibitor of g protein-coupled receptor kinase 2 and increases myocardial contractility. ACS Chem. Biol. 7, 1830–1839 (2012).

70. Sun, X. et al. Paroxetine Attenuates Cardiac Hypertrophy Via Blocking GRK2 and ADRB1 Interaction in Hypertension. J. Am. Heart Assoc. Cardiovasc. Cerebrovasc. Dis. 10, e016364 (2020).

71. Bouley, R. et al. Structural Determinants Influencing the Potency and Selectivity of Indazole-Paroxetine Hybrid G Protein-Coupled Receptor Kinase 2 Inhibitors. Mol. Pharmacol. 92, 707–717 (2017).

72. Ferrero, K. M. & Koch, W. J. GRK2 in cardiovascular disease and its potential as a therapeutic target. J. Mol. Cell. Cardiol. 172, 14–23 (2022).

73. Ren, R., Oakley, R. H., Cruz-Topete, D. & Cidlowski, J. A. Dual Role for Glucocorticoids in Cardiomyocyte Hypertrophy and Apoptosis. Endocrinology 153, 5346–5360 (2012).

74. Severinova, E. et al. Glucocorticoid Receptor-Binding and Transcriptome Signature in Cardiomyocytes. J. Am. Heart Assoc. 8, e011484 (2019).

75. Oakley, R. H. et al. Essential role of stress hormone signaling in cardiomyocytes for the prevention of heart disease. Proc. Natl. Acad. Sci. U. S. A. 110, 17035–17040 (2013).

76. Pianca, N. et al. Glucocorticoid receptor antagonization propels endogenous cardiomyocyte proliferation and cardiac regeneration. Nat. Cardiovasc. Res. 1, 617–633 (2022).

