## Supplementary Materials for "Logic-based machine learning predicts how escitalopram attenuates cardiomyocyte hypertrophy"

### SUPPLEMENTARY METHODS

#### Logic-based network model of cardiomyocyte hypertrophy signaling

We used a published logic-based differential equation model of the cardiomyocyte hypertrophy signaling network(Ryall et al., 2012). Netflux software (bioRxiv 2024.01.11.575227) was used to construct the network model from a literature-based network of proteins and genes previously associated with cardiomyocyte hypertrophy. This network model is composed of 107 nodes (proteins or mRNAs) and 193 edges, with 17 biochemical inputs including a mechanical stretch input. Outputs include hypertrophic markers, beta myosin heavy chain (βMHC), atrial natriuretic hormone (ANP), brain natriuretic peptide (BNP) and cell area. As in previous studies, biochemical or stretch stimulation was simulated by setting input reaction weights to a value of 0.1, representing 10% of saturating activity(Khalilimeybodi et al., 2020). Examining 450 experiments across the literature, this model had a 77% accuracy(Khalilimeybodi et al., 2020). Further, the model correctly predicted the anti-hypertrophic effect of 28 out of 32 drug responses from our previous experimental screen(Reid et al., 2016), further examined by LogiRx below.

#### Linking drug targets to logic-based signaling network using LogiRx

LogiRx uses mechanistic machine learning to map pathways from drugs into a validated logic-based network model. We first identified hits from a previous screen for compounds that inhibit cardiomyocyte hypertrophy(Reid et al., 2016). Protein targets for these active compounds were identified using the drug-target database DrugBank(Wishart et al., 2018) based on gene name. Directed protein interactions in the Omnipath protein interaction database(Ceccarelli et al., 2020; Türei et al., 2016) originating from the KEGG(Kanehisa & Goto, 2000), Reactome(Haw et al., 2011), and SIGNOR(Perfetto et al., 2016) datasets were used as potential connections from drug targets to the hypertrophy network model. Drug candidates with either no listed protein targets or no targets with directed protein interactions were excluded. LogiRx identified top scoring directed pathways from drug targets to the hypertrophy signaling network nodes through Omnipath using the PathLinker algorithm(Gil et al., 2017; Ritz et al., 2016) in Cytoscape(Shannon et al., 2003). PathLinker computes a ranked list of candidate pathways by searching a directed protein interaction network for all pathways from a set of source nodes to a set of target nodes. PathLinker identifies optimal pathways by minimizing a cost function of the product of edge weights along each candidate pathway from sources to sinks(Gil et al., 2017; Ritz et al., 2016). The optimization finds the k shortest loopless paths using Yen's algorithm modified with a "best search first" heuristic (Yen, 1971). The parameter k was set to 200, which produced pathways of length ≤ 3. For each top scoring candidate, the drug pathway was added automatically to a LogiRx-expanded logic-based network model using Netflux.

**Literature validation of LogiRx-expanded models**

The LogiRx-expanded logic-based differential equation model was simulated to test the functional impact of the drug pathways on cardiomyocyte hypertrophy. Each pathway was individually added to the signaling network model to create expanded logic-based differential equation model variants. Each drug-pathway-expanded model was simulated to predict the semi-quantitative network dynamics in response to drug. To assess the functional validity of the LogiRx-identified drug pathways, we validated their performance against 450 literature experiments previously used to validate the hypertrophy signaling model(Khalilimeybodi et al., 2020).

**Simulating drug mechanisms of action**

Drug mechanisms of action were implemented using logic-based equations as described previously(Eggertsen & Saucerman, 2023; Zeigler et al., 2021), accounting for mechanisms of drug action (agonist or antagonist) and drug binding (competitive or non-competitive). The binding properties determine how the upstream signal is shifted by the drug dose, while the drug action determines whether the target node is upregulated (agonist) or downregulated (antagonist). Each simulation was run first without drug input to establish baseline activity, and then with the appropriate drug to identify steady-state changes in drug response.

**Mechanistic subnetwork analysis**

To identify pathways mediating drug response, we utilized the mechanistic subnetwork method(Eggertsen & Saucerman, 2023; Van de Graaf et al., 2023; Zeigler et al., 2021). We simulated the global response of the network to the drug (normalized dose = 1) in the presence of hypertrophic stimulus (input reaction weight for PE or TGFβ = 0.1 with other input weights = 0.02) to identify "drug-responsive nodes". We then simulated knockdown of each node ($Y_{MAX}$ = 0) and measured that node's influence on drug-induced change in cardiomyocyte hypertrophy, referring to these nodes as "regulating nodes". The intersection of the regulating nodes with responsive nodes results in a mechanistic subnetwork that maps the pathways mediating drug activity. Thus, LogiRx combines drug-target and protein interaction databases, path optimization, and logic-based network simulations to obtain mechanistic subnetworks that explain drug response.

**Experimental validation in cardiomyocytes**

We isolated neonatal cardiomyocytes using the NeoMyts kit from Cellutron as described previously(Eggertsen & Saucerman, 2023). Cardiomyocytes were cultured with serum for 24 hr in 96-well microplates, followed by a 24 hr serum starve. Cardiomyocytes were then treated with one of two hypertrophic stimuli (10 µM phenylephrine, 5 ng/ml transforming growth factor β), 10% FBS (positive control), or serum-free media alone (negative control). At the same time, cells were treated with specified concentrations of the appropriate compound (escitalopram, mifepristone, paroxetine, sarpogrelate hydrochloride, or AL082D06) for 48 hrs.

To prepare for immunofluorescent imaging, cardiomyocytes were first fixed with 4% paraformaldehyde for 20 min, and then permeabilized with 0.1% Triton-X for 15 min. Cardiomyocytes were blocked with 1% bovine serum albumin in PBS for 1 hr, then treated with mouse anti-α-actinin primary antibody (Sigma-Aldrich Cat#A7811, RRID:AB_476766) at a concentration of 1:200 overnight. Cardiomyocytes were blocked with 5% goat serum in PBS for 1 hr, then Alexa Fluor-568-conjugated goat anti-mouse secondary antibody (Thermo Fisher Scientific Cat#A11031, RRID:AB_144696) at a concentration of 1:200 was applied for 1 hr. The cells were stained with DAPI prior to imaging.

High-content imaging was performed on the stained cardiomyocytes using an Operetta CLS High Content Analysis System. These images were processed using CellProfiler(Stirling et al., 2021) using a cellular segmentation algorithm developed previously and validated to within 5% of two independent manual segmentations(Bass et al., 2012). Median cell area was used as a representative measure of the cell population in each well, and cells with undetectable cytoplasm were not counted.

**Statistics**

All data are presented as the mean ± SEM. Analysis of experimental conditions considered two distinct cardiomyocyte isolations and multiple conditions, with multiple wells in each experiment. For this reason, a two-way ANOVA followed by Dunnett's test for multiple comparisons was selected for statistical calculation. In vivo data were analyzed by ordinary one-way ANOVA followed by Tukey's multiple comparisons test. GraphPad Prism 9 was used for statistical analysis. For all analyses, a p-value <0.05 was considered statistically significant. Code for LogiRx, the expanded hypertrophy network, and data analyses are on Github at: https://github.com/saucermanlab/Eggertsen_LogiRx.

**SUPPLEMENTARY TABLE**

**Supplementary Table S1.** Limitations and potential improvements to resources used for LogiRx that affect its performance.

| Limitation | Potential Improvements |
|---|---|
| **Incomplete Mapping** | |
| 28 of 62 drugs from Reid et al. 2016 mapped to protein targets in DrugBank/Omnipath | More comprehensive experimental characterization of drug targets |
| 11 of 28 drugs with known protein targets mapped to the hypertrophy network via Pathlinker/Omnipath | More comprehensive experimental characterization of protein interactions<br>Greater coverage of directed protein interactions in databases<br>Identify hypertrophic pathways that do not involve reactions or nodes in the existing hypertrophy network model |
| 76% of inferred drug pathways work via newly inferred intermediate proteins | Consider pathways of length >2 to capture even more drug pathways |
| **Incorrect Model Predictions** | |
| 8 of 11 drugs were predicted to regulate hypertrophy with validation accuracy >78% | Experiments to identify crosstalk between inferred drug pathways and those in network model |
| 7 of 8 drugs were predicted to inhibit PE-induced hypertrophy | For drugs with multiple targets (8 targets modeled for bromocriptine), experimentally characterize differing weights for each target |
| 3 of 7 drugs (escitalopram, tirbanibulin, mifepristone) predicted to inhibit hypertrophy were validated in a PE stimulation context | Inclusion of reactions or stronger reaction weights not tuned for the cell type; decrease experimental variability or increase statistical power; Optimize environmental variables such as serum starve, matrix content/stiffness, dependence on species and developmental stage |
| 2 of 3 drugs (escitalopram, mifepristone) that inhibited hypertrophy in a PE context also inhibited TGFβ-induced hypertrophy | More detailed modeling of context-dependent signaling across biochemical stimuli |
| Original hypertrophy model exhibited 77% accuracy against 450 experiments | Sensitivity analysis to guide additional experiments and model revision |

**SUPPLEMENTARY FIGURES**

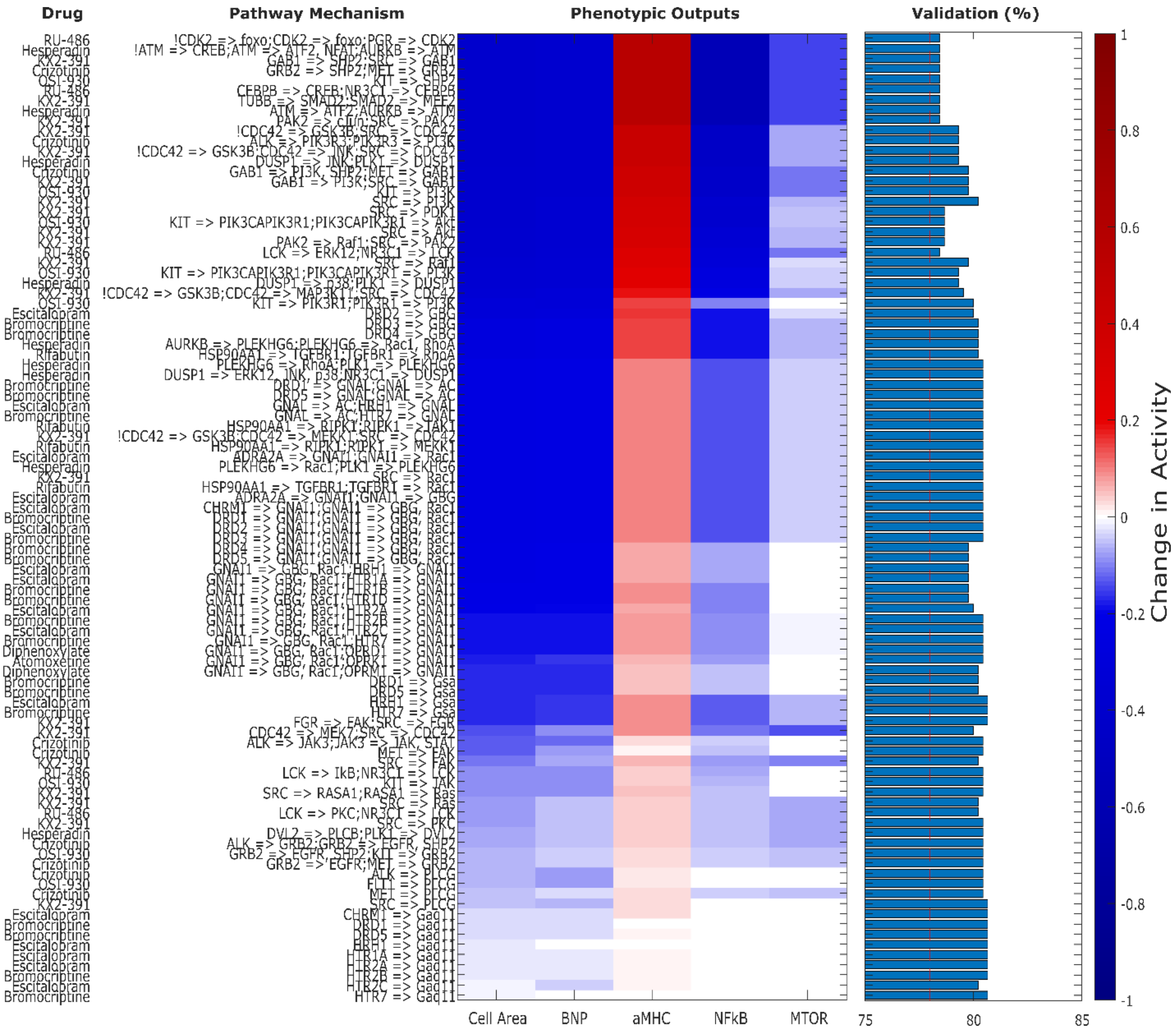


**Supplementary Figure S1. 11 drugs and 95 corresponding drug pathways that map to the hypertrophy network and are predicted to attenuate cardiomyocyte cell area.** Each row depicts an expansion of the hypertrophy network, simulated for their effect on five hypertrophic outputs, as well as validated against 450 experiments from prior literature. This is an annotated version of **Figure 2A**.

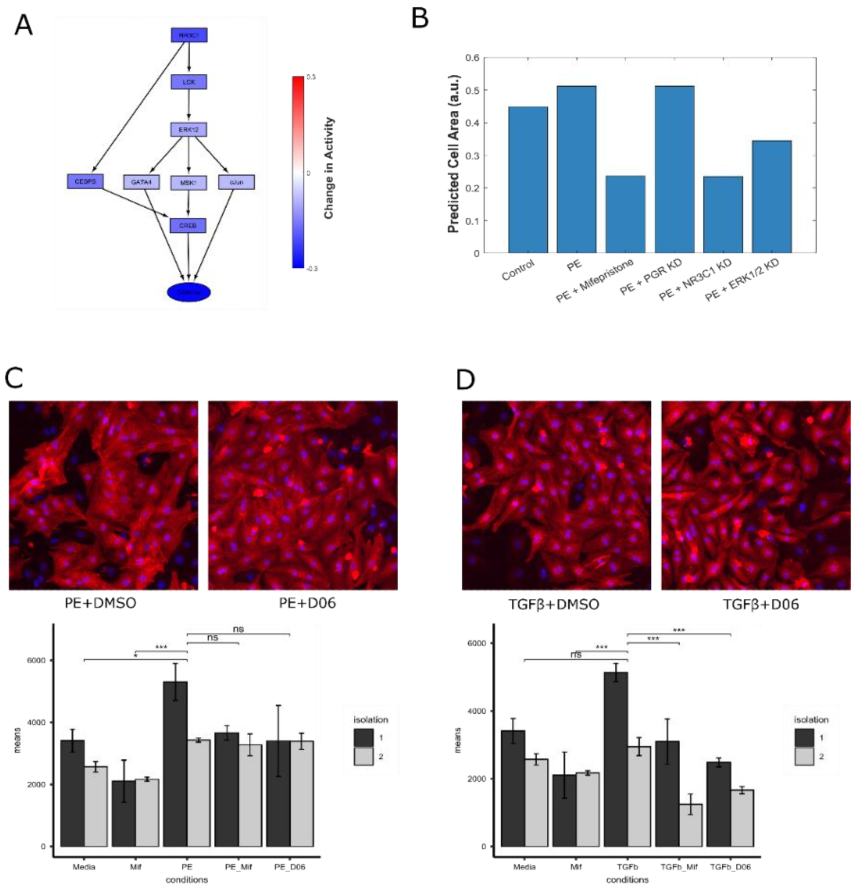


**Supplementary Figure S2. Mifepristone prevents cardiomyocyte hypertrophy through glucocorticoid receptor inhibition.** A) Mechanistic subnetwork analysis predicts that mifepristone suppresses cardiomyocyte hypertrophy through the glucocorticoid receptor via LCK and CEBPβ. Prior studies have shown that the glucocorticoids and the glucocorticoid receptor may be either hypertrophic or antihypertrophic(Ren et al., 2012), and it is not clear what mechanisms control these divergent effects. The role of LCK has not been described in hypertrophy, however it is known to mediate cardioprotection from ischemic injury(Ping et al., 2002). B) Simulations of PE-induced hypertrophy and modulation by mifepristone, progesterone inhibition, glucocorticoid receptor inhibition, or ERK ½ inhibition. Glucocorticoid receptor inhibition by AL082D06 (20 µM) significantly prevents C) PE- and D) TGFβ-induced cardiomyocyte hypertrophy 48 hours post treatment, as shown by representative images and quantification of automated segmentation. CEBPβ has been previously been shown to regulate cardiac hypertrophy(Redondo-Angulo et al., 2016; Zou et al., 2014).

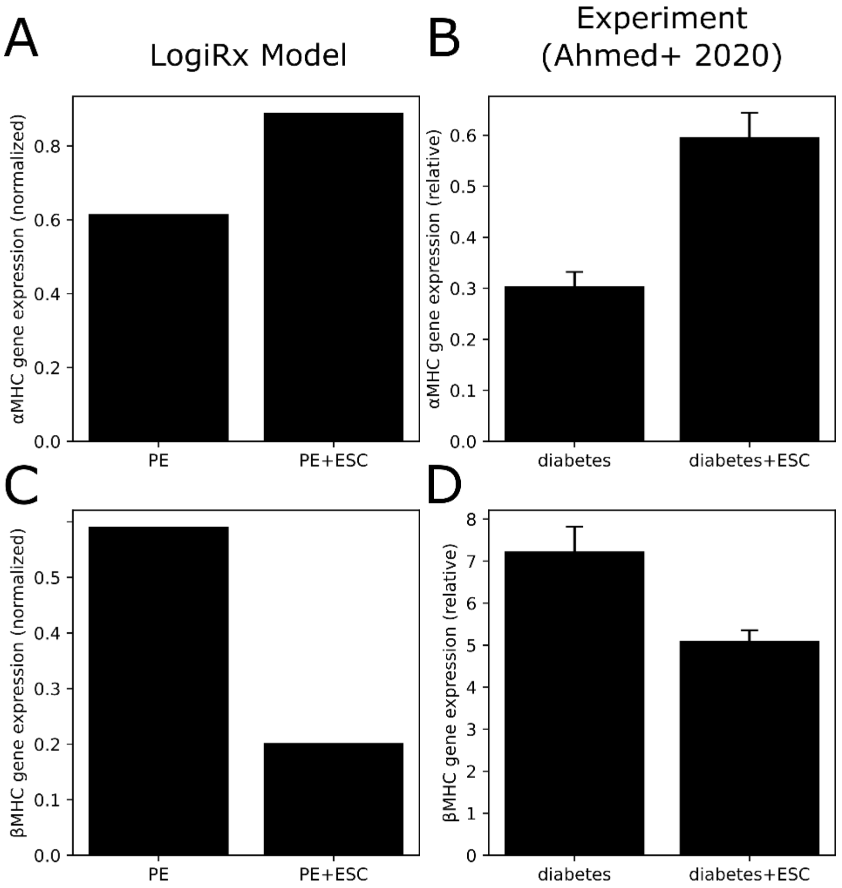


**Supplementary Figure S3. Validation of predicted cardiac gene expression in response to escitalopram.** A) Predicted αMHC gene expression by the LogiRx network model under PE treatment, with and without escitalopram (ESC). B) Experimentally measured αMHC gene expression in rat hearts from a diabetes model (high-fat high-fructose diet with streptozotocin injections), with and without escitalopram. Experimental data from (Ahmed et al., 2020). C) Predicted βMHC gene expression by the LogiRx network model under PE treatment, with and without escitalopram (ESC). B) Experimentally measured βMHC gene expression in rat hearts from a diabetes model, with and without escitalopram. Experimental data from (Ahmed et al., 2020).

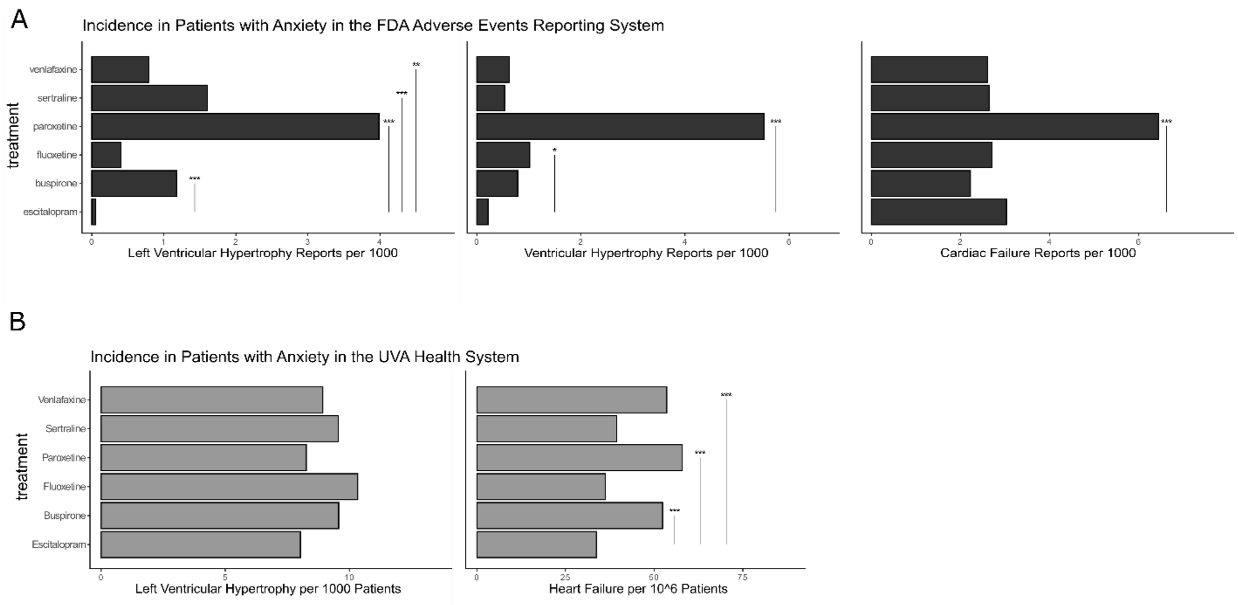


**Supplementary Figure S4. Left ventricular hypertrophy is less frequently reported among anxiety patients taking escitalopram.** A) Reports of left ventricular hypertrophy (left), ventricular hypertrophy (center), or cardiac failure (right) per 1000 patients are shown for anxiety patients taking either escitalopram or another anxiolytic (buspirone, fluoxetine, paroxetine, sertraline or venlafaxine). B) Diagnoses of left ventricular hypertrophy per 1000 (left) or heart failure per 1 million (right) patients are shown for anxiety patients taking either escitalopram or another anxiolytic.

### SUPPLEMENTARY REFERENCES


Ahmed, L. A., Shiha, N. A., & Attia, A. S. (2020). Escitalopram Ameliorates Cardiomyopathy in Type 2 Diabetic Rats via Modulation of Receptor for Advanced Glycation End Products and Its Downstream Signaling Cascades. *Frontiers in Pharmacology*, *11*. https://doi.org/10.3389/fphar.2020.579206

Bass, G. T., Ryall, K. A., Katikapalli, A., Taylor, B. E., Dang, S. T., Acton, S. T., & Saucerman, J. J. (2012). Automated image analysis identifies signaling pathways regulating distinct signatures of cardiac myocyte hypertrophy. *Journal of Molecular and Cellular Cardiology*, *52*(5), 923–930. https://doi.org/10.1016/j.yjmcc.2011.11.009

Ceccarelli, F., Turei, D., Gabor, A., & Saez-Rodriguez, J. (2020). Bringing data from curated pathway resources to Cytoscape with OmniPath. *Bioinformatics*, *36*(8), 2632–2633. https://doi.org/10.1093/bioinformatics/btz968

Eggertsen, T. G., & Saucerman, J. J. (2023). Virtual drug screen reveals context-dependent inhibition of cardiomyocyte hypertrophy. *British Journal of Pharmacology*, *180*(21), 2721–2735. https://doi.org/10.1111/bph.16163

Gil, D. P., Law, J. N., & Murali, T. M. (2017). The PathLinker app: Connect the dots in protein interaction networks. *F1000Research*, *6*, 58. https://doi.org/10.12688/f1000research.9909.1

Haw, R., Hermjakob, H., D'Eustachio, P., & Stein, L. (2011). Reactome pathway analysis to enrich biological discovery in proteomics data sets. *PROTEOMICS*, *11*(18), 3598–3613. https://doi.org/10.1002/pmic.201100066

Kanehisa, M., & Goto, S. (2000). KEGG: Kyoto Encyclopedia of Genes and Genomes. *Nucleic Acids Research*, *28*(1), 27–30. https://doi.org/10.1093/nar/28.1.27

Khalilimeybodi, A., Paap, A. M., Christiansen, S. L. M., & Saucerman, J. J. (2020). Context-specific network modeling identifies new crosstalk in β-adrenergic cardiac hypertrophy. *PLoS Computational Biology*, *16*(12). https://doi.org/10.1371/journal.pcbi.1008490

Perfetto, L., Briganti, L., Calderone, A., Cerquone Perpetuini, A., Iannuccelli, M., Langone, F., Licata, L., Marinkovic, M., Mattioni, A., Pavlidou, T., Peluso, D., Petrilli, L. L., Pirrò, S., Posca, D., Santonico, E., Silvestri, A., Spada, F., Castagnoli, L., & Cesareni, G. (2016). SIGNOR: A database of causal relationships between biological entities. *Nucleic Acids Research*, *44*(D1), Article D1. https://doi.org/10.1093/nar/gkv1048

Ping, P., Song, C., Zhang, J., Guo, Y., Cao, X., Li, R. C. X., Wu, W., Vondriska, T. M., Pass, J. M., Tang, X.-L., Pierce, W. M., & Bolli, R. (2002). Formation of protein kinase Cε-Lck signaling modules confers cardioprotection. *The Journal of Clinical Investigation*, *109*(4), 499–507. https://doi.org/10.1172/JCI13200

Redondo-Angulo, I., Mas-Stachurska, A., Sitges, M., Giralt, M., Villarroya, F., & Planavila, A. (2016). C/EBPβ is required in pregnancy-induced cardiac hypertrophy. *International Journal of Cardiology*, *202*, 819–828. https://doi.org/10.1016/j.ijcard.2015.10.005

Reid, B. G., Stratton, M. S., Bowers, S., Cavasin, M. A., Demos-Davies, K. M., Susano, I., & McKinsey, T. A. (2016). Discovery of novel small molecule inhibitors of cardiac hypertrophy using high throughput, high content imaging. *Journal of Molecular and Cellular Cardiology*, *97*, 106–113. https://doi.org/10.1016/j.yjmcc.2016.04.015

Ren, R., Oakley, R. H., Cruz-Topete, D., & Cidlowski, J. A. (2012). Dual Role for Glucocorticoids in Cardiomyocyte Hypertrophy and Apoptosis. *Endocrinology*, *153*(11), 5346–5360. https://doi.org/10.1210/en.2012-1563

Ritz, A., Poirel, C. L., Tegge, A. N., Sharp, N., Simmons, K., Powell, A., Kale, S. D., & Murali, T. M. (2016). Pathways on demand: Automated reconstruction of human signaling networks. *NPJ Systems Biology and Applications*, *2*, 16002. https://doi.org/10.1038/npjsba.2016.2

Ryall, K., Holland, D., Delaney, K., Kraeutler, M., Parker, A., & Saucerman, J. (2012). Network reconstruction and systems analysis of cardiac myocyte hypertrophy signaling. *The Journal of Biological Chemistry*, *287*(50), Article 50. https://doi.org/10.1074/jbc.m112.382937

Sarangdhar, M., Tabar, S., Schmidt, C., Kushwaha, A., Shah, K., Dahlquist, J. E., Jegga, A. G., & Aronow, B. J. (2016). Data mining differential clinical outcomes associated with drug regimens using adverse event reporting data. *Nature Biotechnology*, *34*(7), 697–700. https://doi.org/10.1038/nbt.3623

Shannon, P., Markiel, A., Ozier, O., Baliga, N. S., Wang, J. T., Ramage, D., Amin, N., Schwikowski, B., & Ideker, T. (2003). Cytoscape: A Software Environment for Integrated Models of Biomolecular Interaction Networks. *Genome Research*, *13*(11), 2498–2504. https://doi.org/10.1101/gr.1239303

Stirling, D. R., Swain-Bowden, M. J., Lucas, A. M., Carpenter, A. E., Cimini, B. A., & Goodman, A. (2021). CellProfiler 4: Improvements in speed, utility and usability. *BMC Bioinformatics*, *22*(1), 433. https://doi.org/10.1186/s12859-021-04344-9

Türei, D., Korcsmáros, T., & Saez-Rodriguez, J. (2016). OmniPath: Guidelines and gateway for literature-curated signaling pathway resources. *Nature Methods*, *13*(12), Article 12. https://doi.org/10.1038/nmeth.4077

Van de Graaf, M. W., Eggertsen, T. G., Zeigler, A. C., Tan, P. M., & Saucerman, J. J. (2023). Benchmarking of protein interaction databases for integration with manually reconstructed signalling network models. *The Journal of Physiology*. https://doi.org/10.1113/JP284616

Wishart, D. S., Feunang, Y. D., Guo, A. C., Lo, E. J., Marcu, A., Grant, J. R., Sajed, T., Johnson, D., Li, C., & Sayeeda, Z. (2018). DrugBank 5.0: A major update to the DrugBank database for 2018. *Nucleic Acids Research*, *46*(D1), Article D1.

Yen, J. Y. (1971). Finding the K Shortest Loopless Paths in a Network. *Management Science*, *17*(11), 712–716. https://doi.org/10.1287/mnsc.17.11.712

Zeigler, A. C., Chandrabhatla, A. S., Christiansen, S. L., Nelson, A. R., Holmes, J. W., & Saucerman, J. J. (2021). Network model-based screen for FDA-approved drugs affecting cardiac fibrosis. *CPT: Pharmacometrics & Systems Pharmacology*, *10*(4), 377–388. https://doi.org/10.1002/psp4.12599

Zou, J., Li, H., Chen, X., Zeng, S., Ye, J., Zhou, C., Liu, M., Zhang, L., Yu, N., Gan, X., Zhou, H., Xian, Z., Chen, S., & Liu, P. (2014). C/EBPβ knockdown protects cardiomyocytes from hypertrophy via inhibition of p65-NFκB. *Molecular and Cellular Endocrinology*, *390*(1–2), 18–25. https://doi.org/10.1016/j.mce.2014.03.007
